# Reduced cholecystokinin-expressing interneuron input contributes to disinhibition of the hippocampal CA2 region in a mouse model of temporal lobe epilepsy

**DOI:** 10.1101/2022.11.09.515872

**Authors:** Alexander C. Whitebirch, Anastasia Barnett, Bina Santoro, Helen E. Scharfman, Steven A. Siegelbaum

**Affiliations:** Departments of Neuroscience and Pharmacology, Kavli Institute for Brain Science, Mortimer B. Zuckerman Mind Brain Behavior Institute, Columbia University Irving Medical Center, New York, NY 10027; Departments of Child Psychiatry; Neuroscience and Physiology; Psychiatry, New York University Langone Health, New York, NY 10016; The Nathan S. Kline Institute for Psychiatric Research, Orangeburg, NY 10962

**Keywords:** pilocarpine, temporal lobe epilepsy, mouse, hippocampus, CA2, interneurons, cholecystokinin, parvalbumin, perineuronal net, inhibition

## Abstract

A significant proportion of temporal lobe epilepsy (TLE) patients experience drug-resistant seizures associated with mesial temporal sclerosis, in which there is extensive cell loss in the hippocampal CA1 and CA3 subfields, with a relative sparing of dentate gyrus granule cells and the CA2 pyramidal neurons. A role for CA2 in seizure generation was suggested based on findings of a reduction in synaptic inhibition (Williamson & Spencer, 1994) and the presence of interictal-like spike activity in resected hippocampal tissue from TLE patients (Wittner et al., 2009). We recently found that in the pilocarpine-induced *status epilepticus* mouse model of TLE there was an increase in CA2 intrinsic excitability associated with a loss of CA2 synaptic inhibition. Furthermore, chemogenetic silencing of CA2 significantly reduced seizure frequency, consistent with a role of CA2 in promoting seizure generation and/or propagation (Whitebirch et al., 2022). In the present study we explored the basis of this inhibitory deficit using immunohistochemical and electrophysiological approaches. We report a widespread decrease in the density of pro-cholecystokinin-immunopositive interneurons and a functional impairment of cholecystokinin-expressing interneuron-mediated inhibition of CA2 pyramidal neurons. We also found a decrease in the density of CA2 parvalbumin-immunopositive interneurons and disruption to the pyramidal neuron-associated perisomatic perineuronal net in the CA2 subfield. These data reveal a set of pathological alterations that may disrupt inhibition of CA2 pyramidal neurons and their downstream targets in epileptic mice.

## INTRODUCTION

Temporal lobe epilepsy (TLE) is among the most prevalent neurological disorders and a proportion of people living with TLE experience refractory seizures that are not effectively controlled by any currently available anti-epileptic drugs nor by surgical resection of seizure-generating tissue (Asadi-Pooya, Stewart, Abrams, & Sharan, 2017; Téllez-Zenteno & Hernández-Ronquillo, 2012). Recurring temporal lobe seizures are associated with a characteristic pattern of hippocampal neurodegeneration and synaptic reorganization, termed mesial temporal sclerosis, both in humans and across numerous animal models of TLE. Particularly notable is the relative resilience of the understudied CA2 region (Blümcke et al., 2013; Sastri et al., 2014; Steve, Jirsch, & Gross, 2014; Thom et al., 2010), a narrow area interposed between the larger CA3 and CA1 regions, initially described in 1934 by Lorente de Nó (Lorente De Nó, 1934). Prior investigations in human TLE tissue have revealed numerous alterations in the CA2 region, including sprouting of dentate gyrus mossy fiber axons (Freiman et al., 2021; Wittner et al., 2009), decreases in parvalbumin expression (Andrioli & Arellano, 2007; Wittner et al., 2009), diminished synaptic inhibition (Williamson & Spencer, 1994), and the appearance of spontaneous interictal-like epileptiform discharges (Wittner et al., 2009).

CA2 pyramidal neurons (PNs) are centrally located within the hippocampal network and receive exceptionally strong entorhinal cortex (EC) input, thus acting as the central node in a powerful disynaptic circuit linking the entorhinal cortex directly to CA1 (Chevaleyre & Siegelbaum, 2010; Cui, Gerfen, & Young, 2013; Hitti & Siegelbaum, 2014; Kohara et al., 2014; Leroy et al., 2017; Sun et al., 2017). CA2 PNs also receive direct input from dentate gyrus (DG) granule cell mossy fibers (Kohara et al., 2014; Whitebirch et al., 2022), and CA2 PN local axonal collaterals contribute to an excitatory recurrent network throughout the CA2 and CA3 subfields (Cui et al., 2013; Hitti & Siegelbaum, 2014; Ishizuka, Weber, & Amaral, 1990; Kohara et al., 2014; Li, Somogyi, Ylinen, & Buzsáki, 1994; Okamoto & Ikegaya, 2018; Tamamaki, Abe, & Nojyo, 1988; Whitebirch et al., 2022). Among the hippocampal subfields CA2 contains the greatest density of inhibitory interneurons, including a population of parvalbumin-expressing (PV+) interneurons with distinct anatomical and physiological properties (Botcher, Falck, Thomson, & Mercer, 2014; Mercer, Eastlake, Trigg, & Thomson, 2012; Mercer, Trigg, & Thomson, 2007). CA2 PNs also receive strong inhibitory input from a population of cholecystokinin-expressing (CCK+) interneurons (Botcher et al., 2014; Loisy et al., 2022; Modi et al., 2019).

Accumulating evidence suggests that CA2 functions as a key circuit node regulating hippocampal network activity and excitability. Sharp-wave ripple (SWR) events are preceded by the selective recruitment of a subset of CA2 PNs, and optogenetic activation of CA2 neurons triggers SWR-associated population reactivation (Oliva, Fernández-Ruiz, Buzsáki, & Berényi, 2016; Oliva, Fernández-Ruiz, Leroy, & Siegelbaum, 2020). Modulation of CA2 PNs influences both the frequency of spontaneous SWRs and the power and coherence of gamma oscillations between the hippocampus and the prefrontal cortex (Alexander et al., 2018; Boehringer et al., 2017). Optogenetic inactivation of CA2 diminishes hippocampal network synchrony and degrades the firing precision of CA1 neurons during both SWRs and behavioral states dominated by theta rhythm (He, Boehringer, Huang, Overton, & Polygalov, 2020; MacDonald & Tonegawa, 2021). Most recently, calcium imaging in mice revealed selective recruitment of lateral EC afferents, CA2 PN axons, and CCK+ interneurons in the CA1 subfield during desynchronized network states (Dudok et al., 2021). Together, the unique synaptic connectivity and functional properties of CA2, the resilience of the CA2 subfield to mesial temporal sclerosis, and alterations in CA2 in human TLE tissue suggest that CA2 may have an important role in the epileptic hippocampus.

Indeed, in the pilocarpine-induced *status epilepticus* (PILO-SE) model of TLE in mice we recently found that chemogenetic inhibition of CA2 PNs *in vivo* significantly reduced the frequency of spontaneous recurring convulsive seizures (Whitebirch et al., 2022). Acute hippocampal slice electrophysiology revealed multiple alterations to the cells and circuitry of the CA2 subfield, including a significant impairment to feedforward inhibition that may facilitate the emergence and propagation of epileptiform activity (Whitebirch et al., 2022). Here, we used electrophysiological and immunohistochemical approaches to explore the mechanisms underlying this inhibitory impairment and found that PILO-SE was associated with alterations to interneuronal populations that may compromise inhibitory control of CA2 excitability in epileptic mice.

## RESULTS

### The CA2a and CA1c subfields are resistant to neurodegeneration following PILO-SE

Our goal here was to provide a detailed quantitative investigation into how the various hippocampal cornu ammonis regions (CA1, CA2 and CA3) are affected in the PILO-SE mouse model of TLE, with particular attention to the mechanisms underlying our previously characterized loss of synaptic inhibitory drive onto CA2 pyramidal cells (Whitebirch et al., 2022). In our previous and current study, we characterized the alterations in the electrophysiological properties of CA2 neurons in acute hippocampal slices from PILO-SE mice that were F1 hybrids from a cross between C57BL/6J and 129S1/SvlmJ mouse lines. We found that this genetic background provided an optimal compromise between induction of SE and mouse survival, which was suitable for subsequent electrophysiological analysis. Although we had previously confirmed at a quantitative level that these mice developed spontaneous seizures, here we first report results from a quantitative analysis of seizure properties in these mice to further validate their use as a TLE model. This is of particular importance as the development of spontaneous seizures in the PILO-SE model is known to be quite sensitive to genetic background, and the commonly used C57BL/6J substrain of mice contains a specific genetic mutation that makes them resistant to seizure generation (Bankstahl, Müller, Wilk, Schughart, & Löscher, 2012; Kapur et al., 2020; Löscher, Ferland, & Ferraro, 2017).

Experiments were performed approximately six weeks following PILO-SE induction (Figure 1A) at a time when spontaneous seizures have generally started to appear (Jain, LaFrancois, Botterill, Alcantara-Gonzalez, & Scharfman, 2019; Mazzuferi, Kumar, Rospo, & Kaminski, 2012; Pitsch et al., 2017). Pilocarpine treatment led to SE in 39.9% of mice in a large cohort of animals (Figure 1B). We captured spontaneous seizures in eight mice using continuous video-EEG recordings and subdural screw electrodes, with recordings taking place at least four weeks after SE (Figure 1C; see Methods). Video-EEG recordings confirmed that SE reliably led to spontaneous recurring convulsive seizures in the weeks following pilocarpine treatment (8/8 mice examined). As previously reported for both human TLE and the PILO-SE model (Baud, Ghestem, Benoliel, Becker, & Bernard, 2019; Baud et al., 2018; Pitsch et al., 2017; Whitebirch et al., 2022), spontaneous seizures tended to occur in a clustered pattern across weeks of recording (Figure 1D). Seizures were large amplitude and rhythmic events highly synchronized across all four electrodes (Figure 1E). Qualitative examination of the time-frequency representation of the spontaneous seizures revealed an increase in the relative power of frequencies between approximately 10 – 50 Hz (Figure 1F).

**Figure 1.**
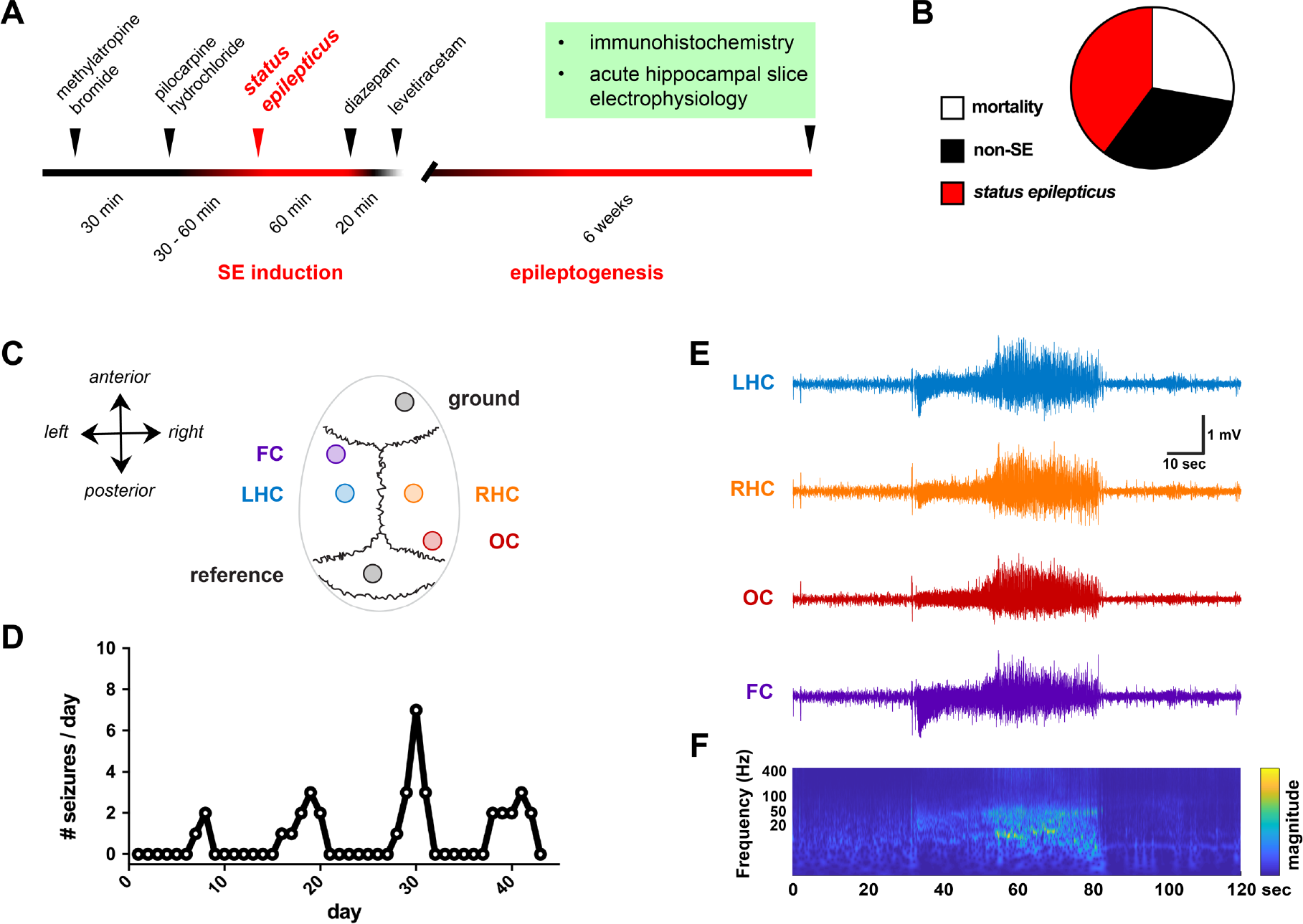
PILO-SE mice experience recurring spontaneous seizures. **(A)** Experimental timeline, in which adult mice are administered pilocarpine hydrochloride (PILO) to induce acute status epilepticus. Immunohistochemistry and *in vitro* electrophysiology are performed approximately 6 weeks following PILO-SE, when mice exhibit spontaneous recurring seizures. **(B)** PILO treatment results in acute mortality (114 of 411 mice, 27.7%), minor seizure activity without status epilepticus (133 of 411 mice, 32.4%), or status epilepticus (164 of 411 mice, 39.9%). **(C)** Seizures were captured through continuous video-EEG recordings, with four screw electrodes positioned over the frontal cortex (FC), left and right hippocampi (LHC and RHC), and occipital cortex (OC). **(D)** Seizure occurrence frequency for one mouse, illustrating the typical clustered pattern of spontaneous recurring seizures. **(E)** A representative seizure, with highly synchronized and large amplitude activity visible across all four electrodes. **(F)** A time- frequency representation of the seizure in panel E, illustrating the increase in power across approximately 1 – 50 Hz.

As previous studies characterizing cell loss in TLE and its rodent models relied solely on anatomical position to distinguish CA2 from its neighboring CA1 and CA3 regions, we next carried out the first quantitative analysis of cell survival using molecular markers to identify specifically CA2 pyramidal neurons. Hippocampal sections from PILO-SE mice revealed a characteristic pattern of hippocampal damage, with neuronal depletion most apparent in the CA3 and CA1 subfields (Figure 2A_1_, A_2_, C_1_, C_2_). To quantify the pattern of pyramidal neuron loss across mice, we stained hippocampal sections for the Nissl substance along with CA2 molecular markers, such as Purkinje cell protein 4 (PCP4) and striatal-enriched protein tyrosine phosphatase (STEP) (Hitti & Siegelbaum, 2014; Kohara et al., 2014). We measured both Nissl and CA2 marker fluorescence intensity along the proximodistal axis of the *stratum pyramidale*, normalizing measurements for both fluorescence intensity and proximodistal position (centered on the CA2 subfield, see Methods). We found decreases in normalized Nissl fluorescence in CA1 and CA3, with the most dramatic decrease in CA3a (Figure 2E). In particular, we noted a stark contrast in the vulnerability of CA3 compared to CA2 PNs, with a consistent decrease in Nissl intensity in the CA3a subregion and resilience in the distal CA2 subregion closest to CA1 (designated CA2a, see Methods), consistent with our visual inspection of PILO-SE tissue (Figure 2B_1_, B_2_, D_1_, D_2_).

**Figure 2.**
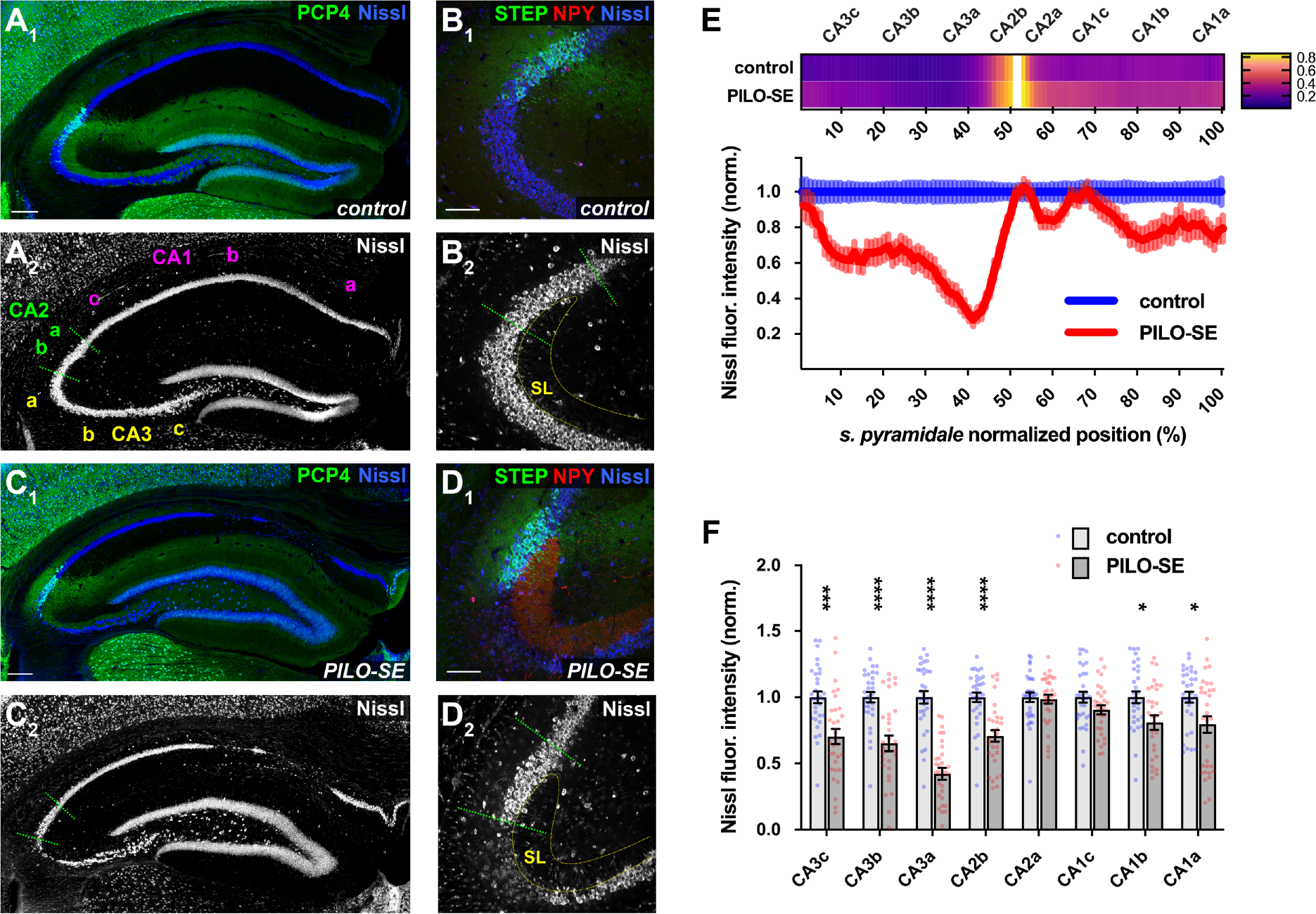
The CA2 subfield was resistant to mesial temporal sclerosis-like neurodegeneration. (A_1_, A_2_) A representative section from a control mouse stained for PCP4 (green) to label CA2 PNs and for Nissl (blue). Scale bar is 200 µm. **(B_1_, B_2_)** CA2 PNs (STEP, green) are located adjacent to CA3a at the distal end of the mossy fiber projection in *stratum lucidum* (SL). Neuronal somata are visualized with a stain for Nissl (blue); a stain for neuropeptide Y (NPY) is shown in red. Scale bar is 100 µm. **(C_1_, C_2_)** Representative mesial temporal sclerosis-like damage in a section from a PILO-SE mouse stained for PCP4 (green) and Nissl (blue). Scale bar is 200 µm. **(D_1_, D_2_)** In sections from PILO-SE mice NPY expression (red) was visible in the mossy fibers. Neuronal somata (Nissl, blue), are absent from CA3a and largely preserved in CA2 (STEP, green). Scale bar is 100 µm. **(E)** Above, a heatmap showing the normalized fluorescence intensity of CA2-specific markers (see methods) along the proximal-distal axis of *stratum pyramidale* in sections from control and PILO-SE mice. Below, normalized Nissl fluorescence intensity across CA3, CA2, and CA1 was decreased significantly in CA3, along with the proximal region of CA2 (CA2b) and the distal portion of CA1 (CA1b, CA1a) (*n* = 45 sections from 31 control mice, 56 sections from 31 PILO-SE mice). **(F)** Measurement of the mean normalized Nissl fluorescence in each subregion confirms a characteristic pattern of neurodegeneration in epileptic mice with mesial temporal sclerosis-like damage, in which CA2a and CA1c are relatively resilient to neurodegeneration.

Measurement of the mean normalized Nissl fluorescence intensity in each subfield (Figure 2F) revealed significant decreases in CA3a (\*\*\*\**P* < 0.0001), CA3b (\*\*\*\**P* < 0.0001), CA3c (\*\*\**P* = 0.0006), CA2b (\*\*\*\**P* < 0.0001), CA1a (\**P* = 0.0326), and CA1b (\**P* = 0.0326) . In contrast, neither CA2a (*P* = 0.7734) nor CA1c (*P* = 0.1505) subregions experienced a significant loss of Nissl signal (two-way ANOVA with Holm-Sidak’s multiple comparisons test; *n* = 45 sections from 31 control mice, 56 sections from 31 PILO-SE mice). Thus our anatomical assessment utilizing established molecular markers of the CA2 region confirmed the previously reported relative resilience of CA2 PNs in the epileptic hippocampus (Blümcke et al., 2013; Mazzuferi et al., 2012; Steve et al., 2014; Winawer et al., 2007). Furthermore, our analysis revealed a differentiation between the proximal and distal section of CA2, with only the latter showing resilience (CA2a). Finally, our measurements revealed a previously unrecognized resilience of the proximal section of CA1 (CA1c) and highlighted the vulnerability of the CA3 region, in particular the CA3a subfield.

### Perisomatic inhibition of CA2 PNs is reduced following pilocarpine administration

Our prior study (Whitebirch et al., 2022) had indicated diminished inhibition of CA2 PNs upon stimulation of either the CA2/CA3 local recurrent collaterals or the DG granule cell mossy fibers following PILO-SE. This loss of inhibition could be explained by a number of mechanisms, including a failure of excitatory axons to sufficiently activate their target interneurons or a reduction in GABA release from recruited interneurons. To explore in more depth the basis of this inhibitory deficit, we performed whole-cell recordings from CA2 PNs in slices from control and PILO-SE mice (approximately six weeks after PILO treatment) in the presence of 50 µM D-AP5 and 25 µM CNQX, antagonists of NMDA and AMPA receptors, respectively. Under these conditions electrical stimulation relatively close to the CA2 subfield directly evoked monosynaptic GABAergic inhibitory postsynaptic potentials (IPSPs) (Figure 3A, B). Electrical stimulation was applied using stimulating electrodes in *stratum pyramidale* (SP), *stratum radiatum* (SR), or *stratum lacunosum moleculare* (SLM) to activate interneuronal axons targeting the perisomatic region (such as those originating from PV+ or CCK+ basket cells), the proximal apical dendrites (where local excitatory axons establish recurrent connections), or the distal apical dendrites (where EC afferents innervate the CA2 region), respectively.

**Figure 3.**
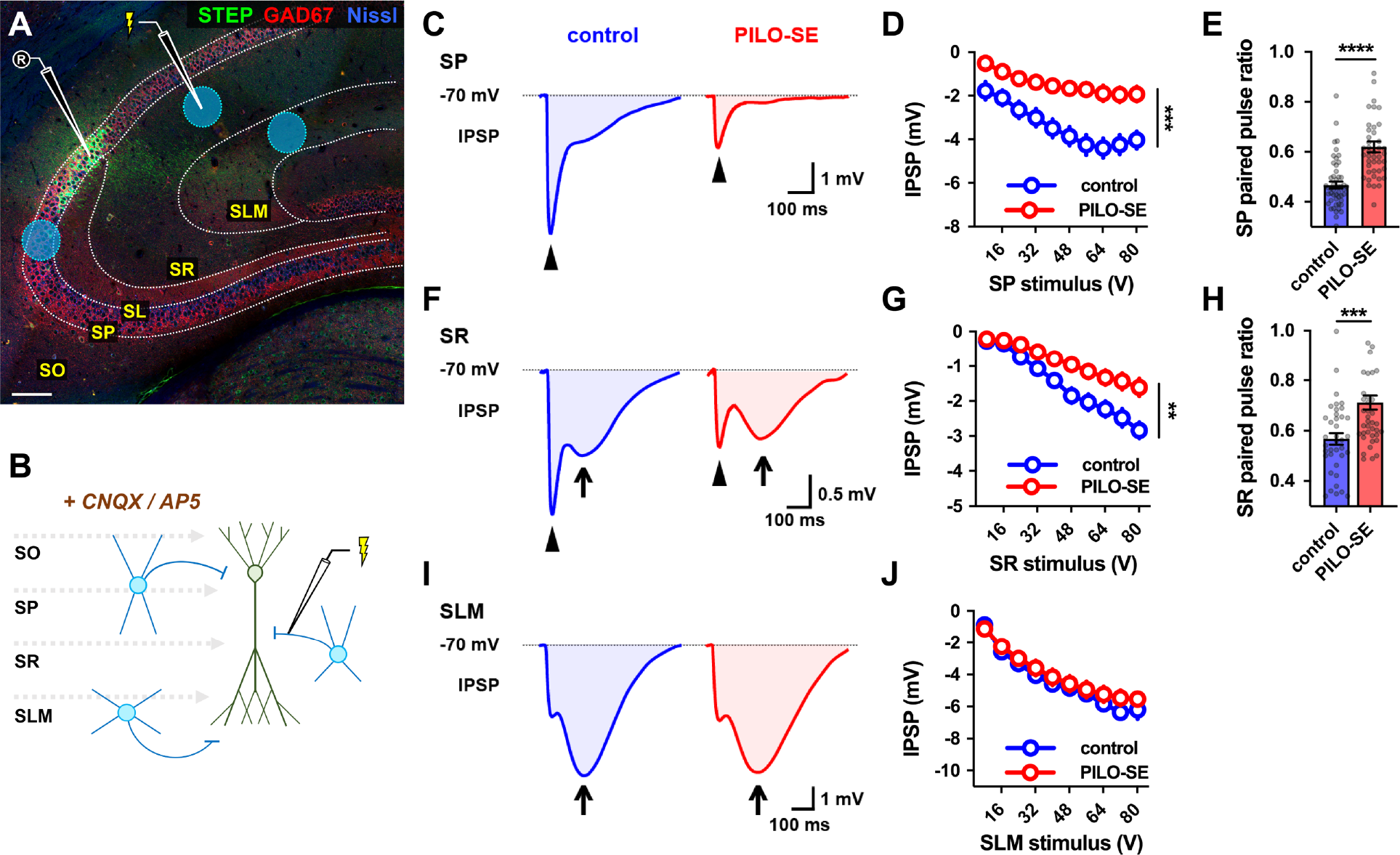
PILO-SE reduced CA2 PN perisomatic and proximal dendritic inhibition, but did not alter inhibition at distal dendrites. **(A)** A representative hippocampal section illustrating the stimulation locations for recruitment of monosynaptic inhibition in *stratum pyramidale* (SP), *stratum radiatum* (SR), and *stratum lacunosum moleculare* (SLM). CNQX and AP5 were added to block excitatory transmission and isolate inhibitory postsynaptic potentials (IPSPs, see Methods). The scale bar is 125 µm. **(B)** A circuit diagram illustrating the experimental configuration: a stimulation pipette located in SP, SR, or SLM evokes monosynaptic inhibition by directly activating local interneurons. **(C)** Representative averaged IPSPs evoked in CA2 PNs by 64V stimulation in SP in slices from control (blue) and PILO-SE (red) mice, with the fast peaks indicated with arrowheads. **(D)** The peak hyperpolarization of the SP stimulation-evoked IPSP was significantly reduced in PILO-SE. **(E)** The paired pulse ratio (PPR) of the IPSP evoked in SP was increased in PILO-SE mice. **(F)** Representative averaged IPSPs evoked by 64V stimulation in SR. The fast peaks are indicated with arrowheads and the slow peaks with arrows. **(G)** The peak hyperpolarization of the SR stimulation-evoked IPSP was significantly reduced in PILO-SE. **(H)** The PPR of the SR stimulation-evoked IPSP was increased in CA2 PNs from PILO-SE mice. **(I)** Representative averaged IPSPs evoked by 64V stimulation in SLM, with the slow peaks indicated by arrows. **(J)** The peak amplitude of the IPSP evoked by SLM stimulation, taking into account both fast and slow phases, was not significantly different between control and PILO-SE.

SP stimulation evoked fast IPSPs in CA2 PNs that were consistent with inhibitory currents mediated by GABA_A_ postsynaptic receptors. On average, PILO-SE caused a reduction in the peak IPSP to 51.7 ± 5.1% of its control value (Mann-Whitney; *\*\*\*\*P <* 0.0001; using a fixed stimulus intensity of 64V, near the peak of the control IPSP input-output curve; n = 36 cells from 23 control mice, 36 cells from 17 PILO-SE mice). Thus, the SP-evoked IPSP input-output curve was significantly reduced in CA2 PNs from PILO-SE mice compared to cells from control animals (Figure 3C, D; mixed-effects model; \*\*\**P* = 0.0001; n = 17 cells from 10 control mice, 20 cells from 7 PILO-SE mice). Pairs of SP stimuli delivered 50 ms apart revealed that PILO-SE caused a significant increase in the paired pulse ratio (PPR), consistent with a decreased probability of GABA release (Figure 3E; Mann-Whitney; \*\*\*\**P* < 0.0001; n = 51 cells from 22 control mice, 44 cells from 18 PILO-SE mice).

Notably, when we delivered stimuli in SR we observed that SR-evoked IPSPs typically exhibited clear fast and slow phases consistent with inhibition mediated by GABA_A_ and GABA_B_ receptors, respectively (Figure 3F). We measured the peak amplitude of the SR-evoked IPSP (taking into account both the fast and slow phases of the response) and found PILO-SE reduced the peak SR-evoked IPSP to 51.1 ± 4.7% of its control value (Mann-Whitney; \*\*\*\**P* < 0.0001; n = 27 cells from 17 control mice, 31 cells from 16 PILO-SE mice), comparable to the reduction in the SP-evoked IPSP. Comparison of the input-output curves for SR-evoked IPSPs revealed a decrease in the magnitude of the IPSP amplitudes (Figure 3F, G; mixed-effects model; \*\**P* = 0.0055; n = 9 cells from 4 control mice, 12 cells from 5 PILO-SE mice). Separate measurement of the peak hyperpolarization during the fast and slow phases of the SR-evoked IPSP revealed a significant decrease to the amplitude of fast inhibition (Kruskal-Wallis with Dunn’s multiple comparisons test; \*\**P* = 0.0034; n = 9 cells from 4 control mice, 12 cells from 5 PILO-SE mice), but no change to the amplitude of slow inhibition (Kruskal-Wallis with Dunn’s multiple comparisons test; *P* = 0.7140; n = 9 cells from 4 control mice, 11 cells from 5 PILO-SE mice). Similar to what we observed with SP stimulation, PILO-SE caused a significant increase in the PPR using paired SR stimuli (Figure 3H; Mann-Whitney; \*\*\**P* = 0.0004; n = 38 cells from 16 control mice, 42 cells from 18 PILO-SE mice). In contrast to the loss of inhibition evoked by stimuli in SP or SR in PILO-SE mice, we found no change in the size of the IPSP evoked by stimulation in the SLM in the epileptic mice compared to control mice, which evoked in both groups biphasic IPSPs dominated by slow inhibition (Figure 3I,J; mixed-effects model; *P* = 0.5548; n = 5 cells from 3 control mice, 16 cells from 7 PILO-SE mice).

Together, these data suggest that the reduction in the IPSP in PILO-SE mice evoked by SP or SR stimulation is caused, at least in part, by a direct monosynaptic effect on inhibitory synaptic transmission. Furthermore, our PPR data indicate that at least part of the decrease in the IPSP is due to a decrease in the probability of transmitter release. Finally, the apparent preferential loss of the fast component of the IPSP evoked by electrical stimulation in SP and SR suggests a selective impairment of fast inhibition mediated by GABA_A_ receptors in CA2 PNs.

### In epileptic mice CA2 parvalbumin-immunopositive interneurons are reduced in density and the CA2 pyramidal neuron-associated perineuronal net is diminished

Perisomatic inhibition in the hippocampus is primarily provided by basket cells that express either cholecystokinin (CCK+) or parvalbumin (PV+) (Dudok et al., 2021; Freund & Katona, 2007). In light of past work demonstrating that CA2 contains a particularly dense population of PV+ interneurons (Botcher et al., 2014), including unique PV+ basket cells (Mercer, Eastlake, et al., 2012; Mercer et al., 2007), we used an immunohistochemical approach to examine PV+ interneurons and perisomatic inhibitory synaptic contacts in the CA2 region (Figure 4). We quantified the density of PV+ interneurons and found a significant decrease in the CA2 SP and SO layers of PILO-SE mice compared to controls (Figure 4C; two-way ANOVA with Holm-Sidak’s multiple comparison test; SP, \**P* = 0.0194; SO, \*\*\**P* = 0.0008; n = 98 sections from 20 control mice, 81 sections from 16 PILO-SE mice). However, the magnitude of the decrease in density of PV+ interneurons in the SP, the location of the basket cells that provide the main inhibitory drive onto CA2 PN soma, was only 12.87 ± 3.48%, which seems unlikely to account for the much larger decrease in the SP-evoked IPSP reported above.

**Figure 4.**
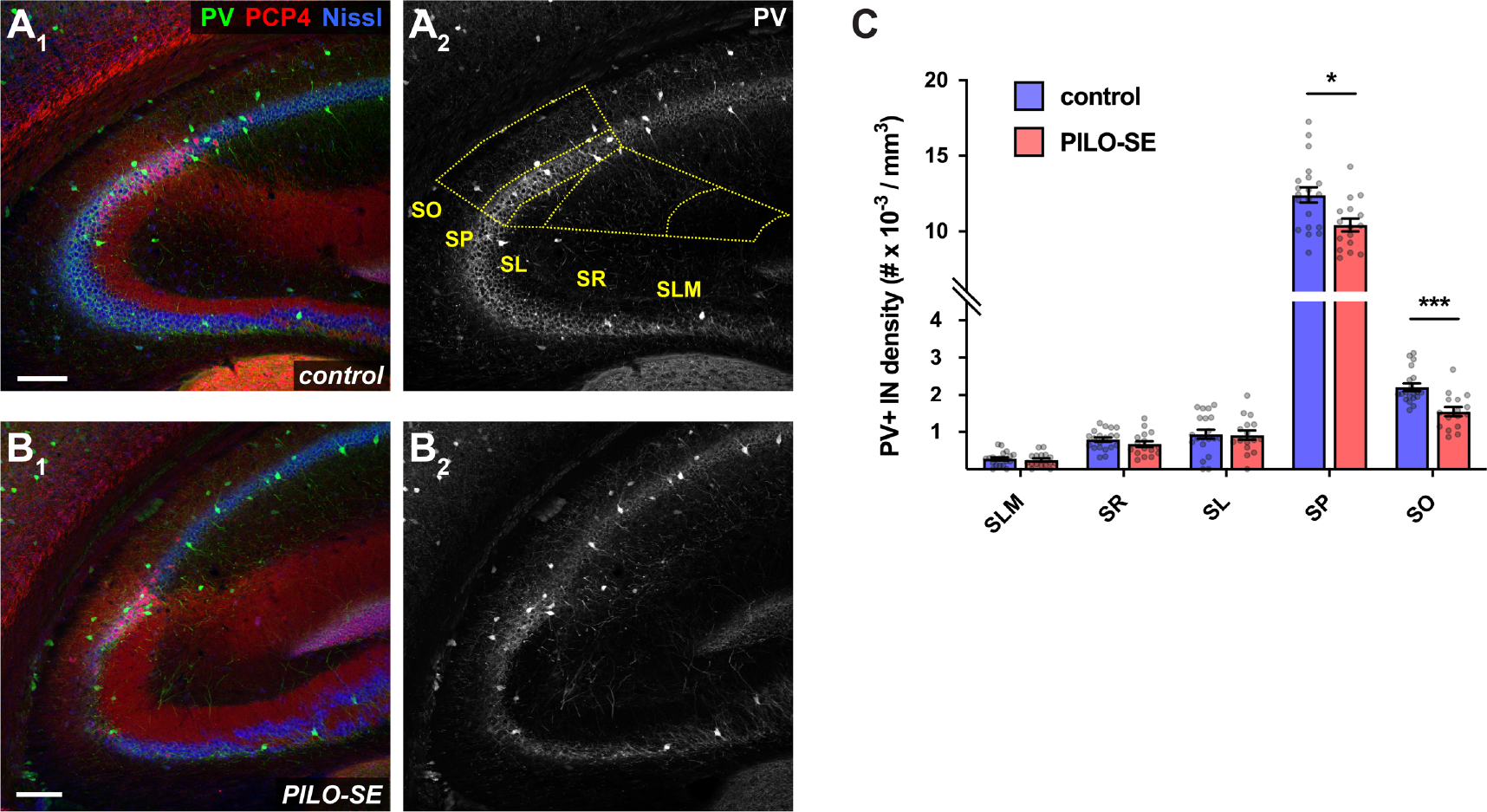
The density of parvalbumin-immunopositive (PV+) interneurons was reduced in the CA2 subfield following PILO-SE. (A_1_ – B_2_) Representative hippocampal sections from control (above) and PILO-SE mice (below), stained for PV (green) to visualize PV+ interneurons, Nissl to label neuronal somata (blue), and PCP4 (red) to delineate the CA2 subfield. The bounds of the CA2 region were drawn across all layers of each hippocampal section (yellow dashed lines in A_2_) and the density of PV+ interneurons was measured for each layer (see methods). Scale bars are 150 µm. **(C)** The density of PV+ INs in CA2 was reduced in the SO and SP layers in sections from PILO-SE mice.

A specialized form of extracellular matrix, the perineuronal net (PNN), provides a structural scaffold that regulates interneuronal excitability and synaptic plasticity (Favuzzi et al., 2017; Slaker, Harkness, & Sorg, 2016; Testa, Prochiantz, & Di Nardo, 2019; Wen, Binder, Ethell, & Razak, 2018). The CA2 subfield exhibits a distinct pattern of PNN expression, with a dense PNN network localized to the perisomatic region of both PV+ interneurons and PNs and a more diffuse PNN extending into the CA2 SO and SR (Noguchi, Matsumoto, Morikawa, Tamura, & Ikegaya, 2017)(Domínguez et al., 2019). Recent evidence has implicated the CA2 PNN in activity-dependent regulation of perisomatic inhibition of CA2 PNs (Carstens, Alexander, & Dudek, 2021; Carstens, Phillips, Pozzo-Miller, Weinberg, & Dudek, 2016; Domínguez et al., 2019). Furthermore, disruption or alteration to the CA2 PNN has been reported in both genetic and chemoconvulsant-based models of epilepsy (Carstens et al., 2021; Dubey et al., 2017; Favuzzi et al., 2017; McRae, Baranov, Rogers, & Porter, 2012; Rankin-gee et al., 2015).

We used the marker *Wisteria floribunda* agglutinin (WFA) to visualize the PNN associated with the CA2 PN perisomatic and proximal dendritic regions, as well as PV+ interneuron soma within the CA2 subfield (Carstens et al., 2016; Carstens et al., 2021; Domínguez et al., 2019). In hippocampal sections we quantified PNN expression in CA2 by measuring the fluorescence intensity of WFA staining in the different layers of each region. We found significant decreases in CA2 WFA intensity (Figure 6A) in tissue from PILO-SE mice as compared to controls in the SP (\*\*\**P* = 0.0001), SR (\*\*\**P* = 0.0002), and SO (\*\**P* = 0.0074) layers (two-way ANOVA with Holm-Sidak’s multiple comparison test; n = 101 sections from 18 control mice, 97 sections from 18 PILO-SE mice). Thus, both the dense meshwork of PNN located around the CA2 PN soma in the SP layer and the diffuse PNN staining in the CA2 SR and SO layers seen in control mice (Figure 5A_1_, A_2_, C_1_, C_2_, E_1_, E_2_) were markedly diminished in PILO-SE mice (Figure 5B_1_, B_2_, D_1_, D_2_, F_1_, F_2_). In contrast, when we examined the pattern of PNN distribution in tissue from PILO-SE mice, we noted that the PNN staining around the soma of PV+ interneurons in CA2 appeared intact, even when it was diminished around CA2 PNs (Figure 5D_1_, D_2_, F_1_, F_2_). Indeed, measurement of WFA fluorescence intensity specifically around PV+ interneurons indicated no differences between control and PILO-SE cells (Figure 6B; Mann-Whitney; *P* = 0.4610; n = 65 cells from 15 control mice, 79 cells from 15 PILO-SE mice), suggesting a selective loss of the pyramidal cell-associated PNN in the CA2 region of epileptic mice. While the mechanisms and functional implications of decreased PNN in CA2 are presently unclear, disruption to the extracellular matrix may be consistent with impairment of inhibitory synaptic function or plasticity in the CA2 region of PILO-SE mice.

**Figure 5.**
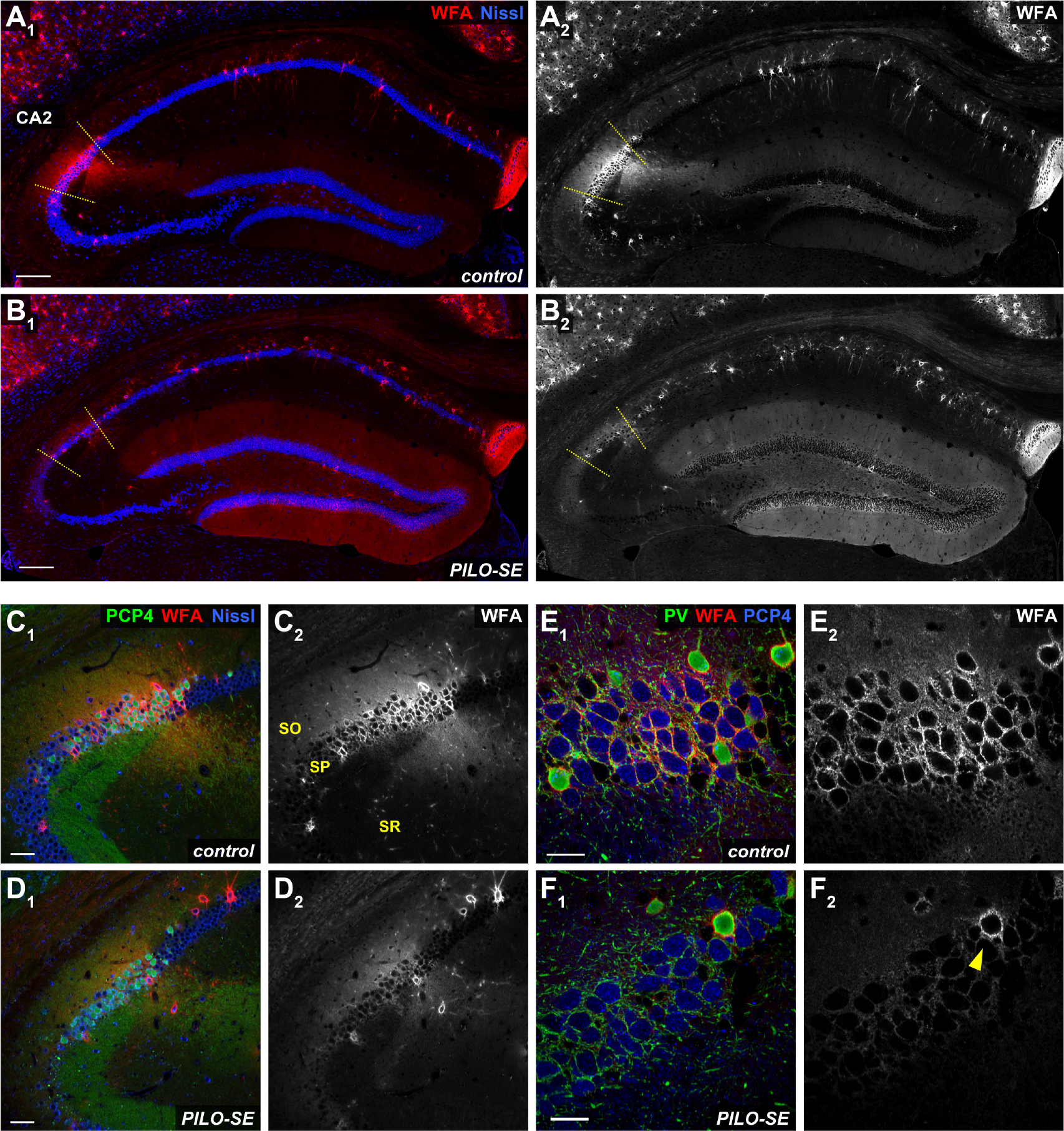
The CA2 pyramidal neuron-associated perineuronal net (PNN) was diminished in pilocarpine-treated mice. **(A_1_, A_2_)** A representative view of hippocampal PNNs, stained with *Wisteria floribunda* agglutinin (WFA) in red. Scale bars are 200 µm. **(B_1_, B_2_)** In PILO-SE mice the intensity of the WFA stain associated with CA2 PNs was diminished while the intensity of the WFA stain in the DG was enhanced. **(C_1_, C_2_)** A representative view of PNNs in CA2. Scale bars are 60 µm. **(D_1_, D_2_)** In PILO-SE mice the intensity of the WFA stain associated with CA2 PNs was diminished. **(E_1_, E_2_)** The perisomatic PNN (stained with WFA in red) surrounds both PV+ interneurons (green) and CA2 PNs (stained with PCP4 in blue). Scale bars are 30 µm. **(F_1_, F_2_)** The PNN was degraded around CA2 PNs but preserved around PV+ interneuron somata in PILO-SE.

**Figure 6.**
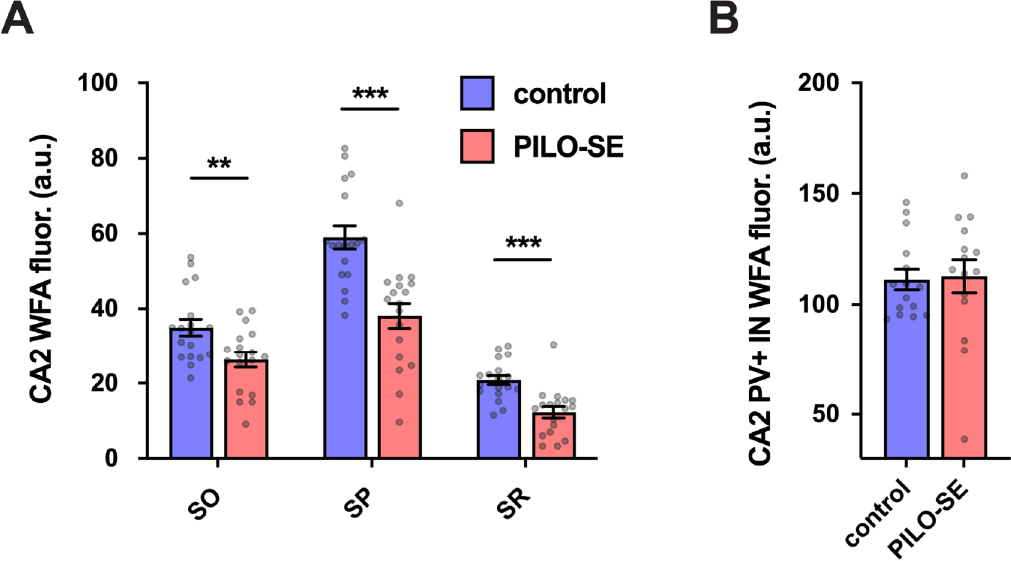
The CA2 pyramidal neuron-associated perineuronal net (PNN) was diminished in pilocarpine-treated mice. **(A)** WFA fluorescence intensity in the *stratum oriens* (SO), *stratum pyramidale* (SP), and *stratum radiatum* (SR) of CA2 was significantly reduced in PILO-SE. **(B)** WFA staining around CA2 PV+ IN somata was not significantly altered in PILO-SE.

### PILO-SE is associated with decreased density and functional impairment of CA2 CCK+ interneurons

Given the discrepancy between the substantial impairment of perisomatic inhibition we observed in CA2 PNs from epileptic mice with the relatively small decrease of parvalbumin-immunopositive somata density, we wondered whether the CCK+ interneurons, which comprise another major population of perisomatic-targeting cells, might be impacted following PILO-SE. We assessed the fate of the CCK+ interneuron population using an antibody for pro-cholecystokinin (pCCK) and found a substantial decrease in the number of pCCK-immunopositive (pCCK+) interneurons in hippocampal sections from PILO-SE mice (Figure 7A_1_ – B_2_). Of particular interest, the magnitude of the decrease in pCCK+ interneuron density was significantly greater than the loss of PV+ IN staining, with an approximately 69.3 ± 4.7% decrease in CA2 pCCK+ positive neurons compared to a 15.9 ± 3.0% decrease in CA2 PV+ interneuron density, when measured across all layers of the hippocampus. This suggests that a disproportionate depletion of CCK+ interneurons in epileptic mice might contribute to the functional loss of inhibition we observed. Measurement of pCCK+ interneuron density in each layer of CA2 revealed significant decreases in pCCK+ interneuron density in all areas except the SLM (Figure 8B; two-way ANOVA with Holm-Sidak’s multiple comparison test; SR, \*\*\*\**P* < 0.0001; SL, \*\*\*\**P* < 0.0001; SP, \*\*\*\**P* < 0.0001; SO, \*\*\*\**P* < 0.0001; n = 78 sections from 15 control mice, 55 sections from 11 PILO-SE mice), a pattern of loss consistent with the layer-specific pattern of IPSP change seen in CA2 PNs from PILO-SE mice (Figure 3).

**Figure 7.**
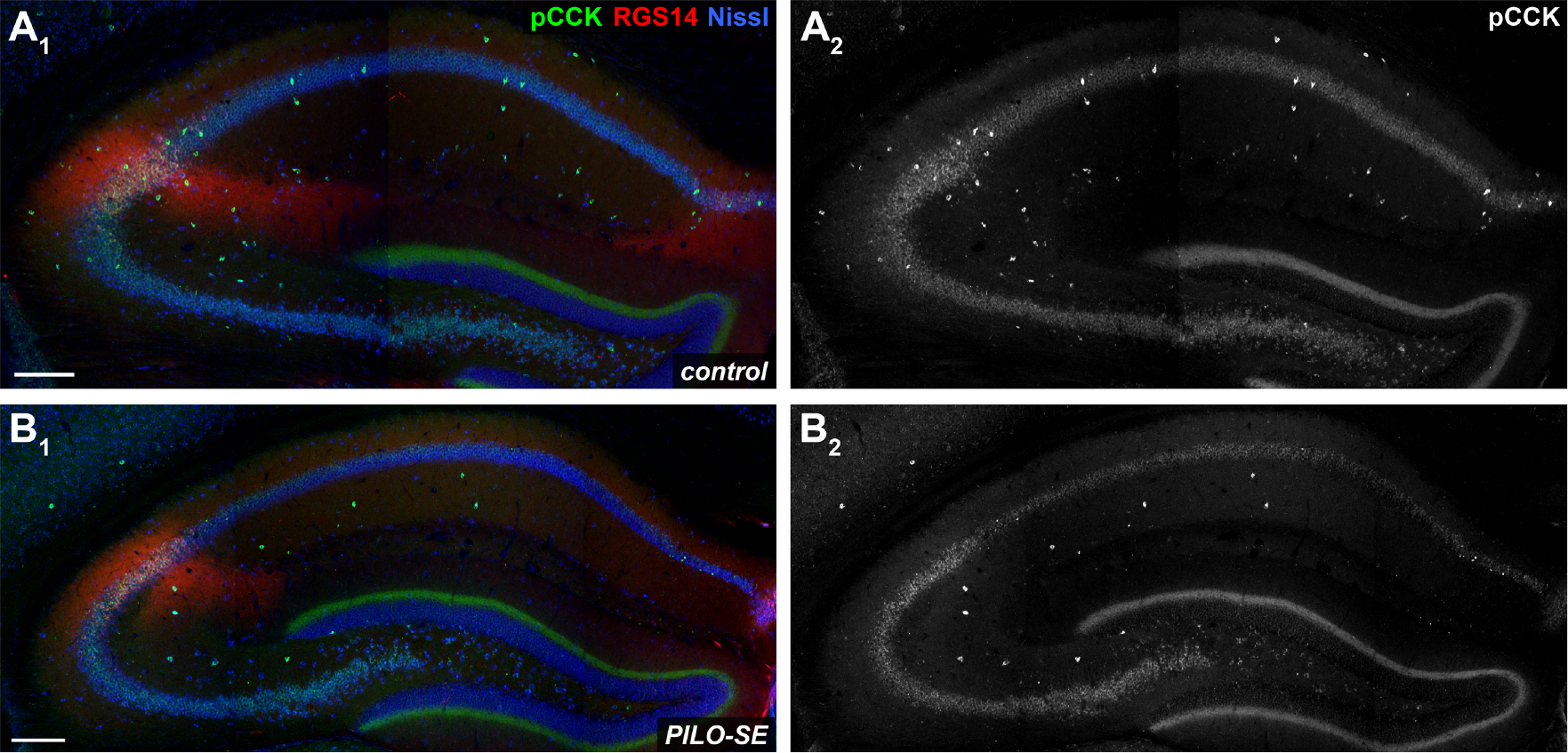
PILO-SE was associated with a widespread decrease of pro-cholecystokinin-immunopositive (pCCK+) interneurons. (A_1_ – B_2_) Representative hippocampal sections from control (A_1_ – A_2_) and PILO-SE (B_1_ – B_2_) mice, stained for pro-cholecystokinin (pCCK, green) to visualize putative cholecystokinin-expressing interneurons, Nissl to label neuronal somata (blue), and RGS14 (red) to delineate the CA2 subfield. Scale bars are 200 µm.

**Figure 8.**
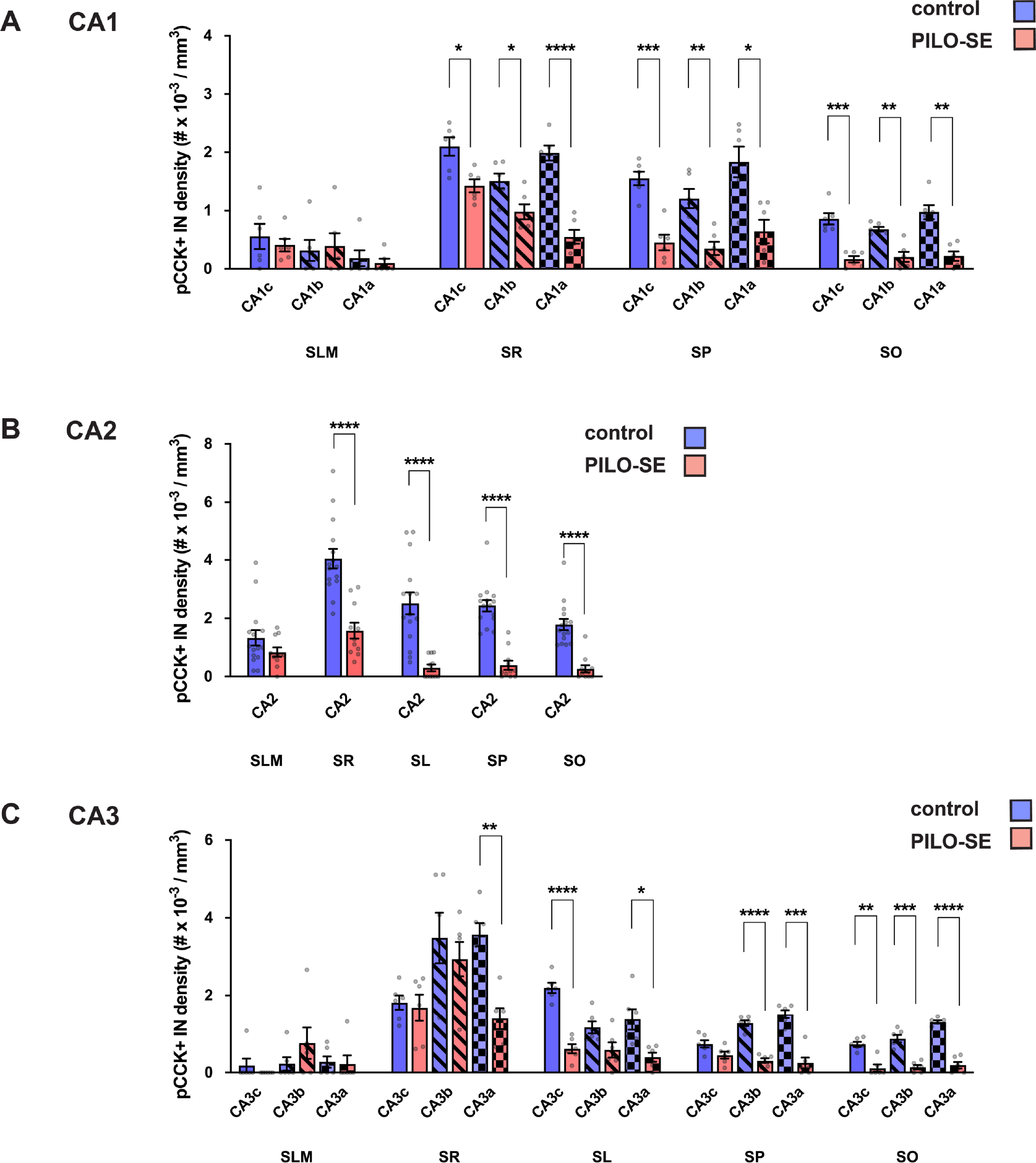
The density of pro-cholecystokinin-immunopositive (pCCK+) interneurons was reduced in CA2 in PILO-SE. The density of pCCK+ interneurons was significantly reduced across all subfields of the hippocampus. **(A)** pCCK+ interneuron density was reduced in CA1 in SO, SP, and SR. **(B)** pCCK+ interneuron density in CA2 was reduced in PILO-SE in SO, SP, SL, and SR. **(C)** pCCK+ interneuron density was reduced in CA3 in SO, SP, SL, and SR, although statistical significance of decrease varied in some layers of CA3b and CA3c, as indicated.

Additionally, we observed a significant and widespread decrease of pCCK+ interneuron density in the other hippocampal subfields (Figure 7). In tissue from PILO-SE mice pCCK+ interneuron density was reduced across the SR, SP and SO layers of CA1c (CA1c SR, \**P* = 0.0133; CA1c SP, \*\*\**P* = 0.0004; CA1c SO), CA1b (\*\*\**P* = 0.0007; CA1b SR, \**P* = 0.0319; CA1b SP, \*\**P* = 0.0073; CA1b SO), and CA1a subfields (\*\**P* = 0.0073; CA1a SR, \*\*\*\**P* < 0.0001; CA1a SP, \**P* = 0.0107; CA1a SO, \*\**P* = 0.0016) when compared to staining in control mice (Figure 8A; two-way ANOVA with Holm-Sidak’s multiple comparison test; n = 33 sections from 6 control mice, 31 sections from 6 PILO-SE mice). Similarly, we found decreases in pCCK+ interneuron density in CA3c (CA3c SR, *P* = 0.7353; CA3c SL, \*\*\*\**P* < 0.0001; CA3c SP, *P* = 0.1436; CA3c SO, \*\**P* = 0.0013), CA3b (CA3b SR, *P* = 0.5034; CA3b SL, *P* = 0.1210; CA3b SP, \*\*\*\**P* < 0.0001; CA3b SO, \*\*\**P* = 0.0006), and CA3a (CA3a SR, \*\**P* = 0.0010; CA3a SL, \**P* = 0.0278; CA3a SP, \*\*\**P* = 0.0002; CA3a SO, \*\*\*\**P* < 0.0001) compared to control animals (Figure 8C; two-way ANOVA with Holm-Sidak’s multiple comparison test; n = 33 sections from 6 control mice, 31 sections from 6 PILO-SE mice). Overall, pCCK+ IN density was reduced across all subregions and layers measured, with the exception of the SR layer of CA3c and CA3b, the SL layer of CA3b, and the SP layer of CA3c.

As the decrease in pCCK+ interneuron density may reflect loss of pCCK protein rather than degeneration of pCCK-expressing interneurons, we next examined whether PILO-SE reduced the IPSP evoked by CCK+ interneurons in CA2 PNs. We took a pharmacological approach based on the distinct neuromodulatory profiles of PV+ and CCK+ interneurons, in which CCK+ interneurons, but not PV+ interneurons, express the cannabinoid type-1 receptor (CB1) that upon activation acts presynaptically to suppress GABA release (Freund & Katona, 2007; Katona et al., 1999). Although CCK+ basket cells are not the only hippocampal interneuronal population that expresses CB1 receptors, by targeting the electrical stimulation to the SP we were able to primarily activate the perisomatic-targeting axons from CCK+ and PV+ basket cells, of which only the former express CB1 receptors (Katona et al., 1999; Wyeth, Zhang, Mody, & Houser, 2010). Because of this anatomical and molecular selectivity, CB1 receptor-mediated depression of perisomatic inhibition has been widely used as a proxy to assess CCK+ basket cell function (Drexel et al., 2017; C. Sun, Sun, Erisir, & Kapur, 2014; Valero et al., 2015; Wyeth et al., 2010). To prevent any modulatory contribution to synaptic transmission of CB1 receptors expressed in excitatory neurons, stimulation was performed in the continual presence of glutamate receptor blockers.

We first used electrical stimulation in the SP layer to evoke direct monosynaptic IPSPs in CA2 PNs. We then applied the CB1 agonist WIN55,212-2 (WIN), which selectively suppresses GABA release from CCK+ interneurons, revealing a WIN-insensitive component of inhibition from PV+ interneurons and other interneuronal populations (including any CCK+ interneuron-mediated inhibition not fully suppressed by WIN) (Figure 9A, B). By measuring the difference between the initial baseline IPSP and the remaining component of the IPSP in the presence of WIN, we obtained the WIN-sensitive component of the IPSP, which provided a measure of the CCK+ interneuron component of the baseline IPSP.

**Figure 9.**
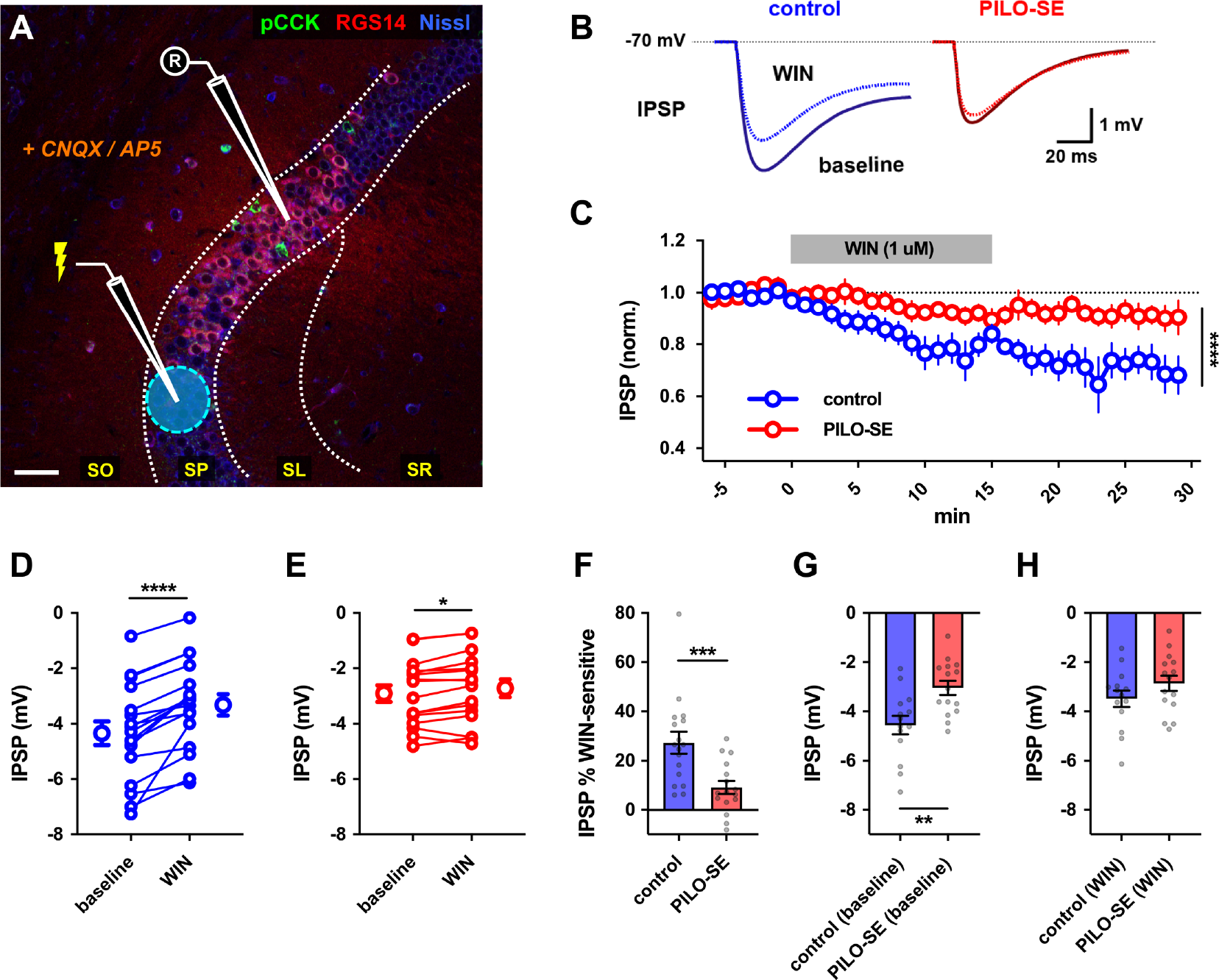
Reduced contribution of CCK+ interneurons to inhibition of CA2 PNs in PILO- SE. **(A)** A representative image illustrating the experimental configuration using electrical stimulation in SP to evoke monosynaptic IPSPs in CA2 PNs (excitation blocked with CNQX and D-APV). Scale bar is 60 µm. **(B)** Representative averaged IPSPs from control (blue) or PILO-SE (red) CA2 PNs before and 11 – 25 min after application of WIN 55,212-2 (WIN, 1 uM). **(C)** Time course of normalized IPSP amplitude following WIN application in CA2 PNs from control and PILO-SE mice.. **(D, E)** IPSPs before and 11 – 25 min after application of WIN in control (D) and PILO-SE (E) mice. Each pair of points is from one cell in separate slices. **(F)** The percent reduction of the IPSP (relative to baseline) in control mice was significantly greater than in PILO-SE mice). **(G)** The baseline (pre-WIN) IPSP amplitude in CA2 PNs from control mice was significantly larger in amplitude than the baseline IPSP amplitude in PILO-SE cells. **(H)** There was no significant difference between the post-WIN IPSP amplitude in CA2 PNs from PILO-SE mice and the post-WIN IPSP amplitude in control cells.

In CA2 PNs from control mice, application of WIN induced a gradual depression of the SP-evoked IPSP that stabilized after approximately 20 mins (Figure 9B, C), resulting in a significant decrease in IPSP amplitude (Figure 9D; Wilcoxon test; \*\*\*\**P* < 0.0001; n = 17 cells from 13 control mice). Consistent with a decrease in pCCK+ interneuron density, the absolute magnitude of the WIN- sensitive component of the IPSP was reduced in PILO-SE mice, from a -1.1 ± 0.2 mV decrease in IPSP size in control cells to a -0.2 ± 0.1 mV decrease in cells from PILO-SE mice (n = 14 cells from 13 control mice, 15 cells from 11 PILO-SE mice). Thus, PILO-SE reduced the magnitude of the WIN-sensitive component of the IPSP by 82.7 ± 6.0 % relative to its value in control animals (Mann-Whitney; \*\*\*\**P* < 0.0001). However, although greatly reduced in amplitude, the effect of WIN to inhibit the IPSP was still significant in PILO-SE mice (Figure 9B, C, E; Wilcoxon test; \**P* = 0.0131; n = 16 cells from 12 PILO-SE mice). Whereas the WIN-sensitive component comprised 27.4 ± 4.4% of the total IPSP in control cells, it comprised only 9.2 ± 2.6% of the IPSP in PILO- SE, a decrease to 33.6% of its value in control animals (Figure 9F; Mann-Whitney; \*\*\**P* = 0.0009; n = 17 cells from 13 control mice, 16 cells from 12 PILO-SE mice).

In principle, the decrease in the WIN-sensitive component of the IPSP in PILO-SE could reflect a loss of sensitivity to CB1 agonist, rather than an actual decrease in the CCK+ interneuron mediated component of the IPSP. The baseline (pre-WIN) IPSP amplitude in CA2 PNs from control mice was significantly larger in amplitude than the baseline IPSP amplitude in PILO-SE cells (Figure 9H; Mann-Whitney; \*\**P* = 0.0043; n = 14 cells from 13 control mice, 15 cells from 11 PILO-SE mice). Importantly, however, we found that the amplitude of the IPSP in control mice in the presence of WIN was identical to that in PILO-SE mice, either in the absence or presence of WIN (Figure 9G, H; Mann-Whitney; control (WIN) versus PILO-SE (baseline), *P* = 0.4509; control (WIN) versus PILO-SE (WIN), *P* = 0.2340; n = 14 cells from 13 control mice, 15 cells from 11 PILO-SE mice). Although the equivalence of the IPSPs could be coincidental, the simplest explanation is that PILO-SE leads to a selective decrease in the size of the CCK+ interneuron mediated IPSP recorded in CA2 PNs. Taken together, these data suggest that a selective loss of CCK+ interneurons from the CA2 subfield contributes to the inhibitory impairment in epileptic mice following PILO-SE.

## DISCUSSION

### Surviving CA2 pyramidal neurons in epileptic mice exhibit impaired perisomatic inhibition

Accumulating evidence indicates that impaired balance and coordination in inhibitory-excitatory networks is a fundamental feature of epilepsy, and prior reports from human TLE tissue have indicated disinhibition of the CA2 subfield (Williamson & Spencer, 1994; Wittner et al., 2009). Our prior study in the PILO-SE mouse model of TLE revealed a widespread loss of inhibition in CA2 circuits. Thus, stimulation of the CA2/CA3 collaterals, CA2 recurrent circuits, or the granule cell mossy fibers evoked significantly smaller inhibitory responses in CA2 PNs from PILO-SE animals (Whitebirch et al., 2022). Furthermore, feedforward inhibitory responses recorded in CA3 and CA1 PNs evoked by selective photostimulation of CA2 axons were also smaller in epileptic mice (Whitebirch et al., 2022). Here we sought to uncover the underlying causes of this inhibitory deficit, reasoning that a decrease in inhibition could result from a number of mechanisms that disrupt distinct components of the feedforward inhibitory circuits, including depletion of specific interneuronal populations or reduced presynaptic GABA release.

When we pharmacologically abolished excitatory neurotransmission and directly evoked monosynaptic inhibition in CA2 PNs, we found a loss of inhibition in slices from epileptic mice that was most apparent in the early phase of the evoked postsynaptic potentials, suggesting a selective impairment in fast GABA_A_-mediated inhibitory currents (Figure 3). We previously observed a similar loss of fast inhibition of CA2 PNs recruited by SR stimulation in the absence of excitatory blockade (Whitebirch et al., 2022). A loss of fast inhibition could result from depletion or dysfunction of interneurons that provide synaptic input to PC somata and proximal dendrites, including (though not limited to) perisomatic-targeting interneurons such as parvalbumin-expressing (PV+) and cholecystokinin-expressing (CCK+) classes of basket cells (Botcher et al., 2014). Our finding of an increased paired-pulse ratio in the PILO-SE mice suggests that a decrease in the probability of GABA release from inhibitory cell presynaptic terminals may contribute to the decreased level of inhibition. It is important to note that our data do not exclude the possibility that an indirect action on disynaptic inhibition resulting from a decrease in the ability of excitatory afferents to effectively recruit interneurons additionally contributes to the impaired feedforward inhibition of CA2 PNs we previously reported (Whitebirch et al., 2022).

Notably, we did not observe a significant reduction in inhibition evoked by stimulation near the PN distal apical dendrites in the SLM. This preservation of the SLM-evoked IPSP, which was characterized by a predominance of slow, likely GABA_B_ receptor-mediated inhibition (Capogna, 2011), may reflect the preferential survival of interneuronal populations that target PN distal apical dendrites such as the neurogliaform cells (Capogna, 2011). Conversely, the clear reduction in SP- and SR-evoked IPSP amplitude may reflect a selective loss or impairment of interneuron populations that target those layers of the hippocampus, including SP-SR interneurons (Mercer, Botcher, Eastlake, & Thomson, 2012), bistratified cells (Mercer et al., 2007), and PV+ or CCK+ basket cells (Mercer, Eastlake, et al., 2012). Although it is possible that some PV+ or CCK+ interneurons may also be recruited by SLM stimulation, relatively few of the somata or dendrites of these interneuron classes are found in the SLM (Botcher et al., 2014).

### Cholecystokinin-expressing interneurons are vulnerable in epileptic mice and their loss may compromise inhibitory control of CA2 excitability

In contrast to the relatively small decrease in PV immunostaining in PILO-SE, we observed a striking loss in staining for pre-cholecystokinin (pCCK). Although we cannot exclude the possibility that pCCK protein decreases while the CCK+ interneurons survive in PILO-SE, we found a large (three-fold), statistically significant decrease in the magnitude of the CCK+ interneuron component of the IPSP, as measured by sensitivity to the CB1 receptor agonist WIN 55,212-2 (WIN), recorded in CA2 PNs (Figure 9). In principle, the reduced effect of WIN to downregulate the IPSP in PILO-SE could potentially result from decreased function or expression of CB1 receptors in CCK+ interneurons (Karlócai et al., 2011; Maglóczky et al., 2010). However, our finding that the magnitude of the CA2 IPSP in the presence of WIN in control animals was similar in size to the CA2 IPSP in the presence of WIN in PILO-SE animals is consistent with a true decrease in the size of the CCK+ interneuron-dependent component of the IPSP in PILO-SE. Conversely, it suggests than any effect of PILO-SE to decrease inhibition mediated by PV+ interneurons is relatively small.

Further support for a decrease in magnitude of the inhibition from CCK+ interneurons comes from a consideration of the effect of PILO-SE on the time course of decay of the inhibitory response recorded in CA2. Although both PV+ and CCK+ basket cells target the perisomatic region, the CCK+ IPSP exhibits a distinctly slower time course of decay, due in part to slow, asynchronous GABA release, which stands in sharp contrast to the rapid and highly synchronous dynamics of PV+ interneuron-mediated inhibition (Basu et al., 2013; Hefft & Jonas, 2005). A selective loss of CCK+ interneurons is consistent with our recent findings that the inhibitory postsynaptic current (IPSC) in CA2 PNs from PILO-SE animals decays more rapidly than the IPSC in control cells (Whitebirch et al., 2022). Thus, the most parsimonious interpretation of our combined immunohistochemical and electrophysiological data is that there is a significant reduction in the contribution of CCK+ interneurons to the IPSP, possibly due to the degeneration of CCK+ interneurons, consistent with several recent investigations in chemoconvulsant-treated animals (Kang et al., 2021; Khan, Shekh-Ahmad, Khalil, Walker, & Ali, 2018; C. Sun et al., 2014; Wyeth et al., 2010). Notably, Wyeth et al., 2010 reported degeneration of CCK-immunopositive and CB1R-immunopositive synaptic boutons in the pyramidal cell layer after PILO-SE in mice, while, in contrast, PV-immunopositive perisomatic boutons were conserved (Wyeth et al., 2010).

In light of the recent evidence linking CB1R-mediated inhibitory plasticity in CA2 CCK+ interneurons to social memory in mice (Loisy et al., 2022), it is intriguing to consider that a loss of CCK+ interneurons in epileptic mice may contribute to social behavioral comorbidities associated with TLE in human patients and animal models of seizures (Ito et al., 2007; Lopes et al., 2016; Okruszek et al., 2017; Seo et al., 2013; Smolensky et al., 2019; Steiger & Jokeit, 2017). Additionally, recent calcium imaging experiments from Dudok et al., 2021 revealed selective recruitment of CCK+ interneurons, lateral EC afferents, and CA2 axons in the CA1 subfield during non-theta, non-SWR desynchronized network activity. These desynchronized network states, which occurred at the transition from locomotion to quiet immobility, were characterized by high levels of CCK+ interneuron activity, widespread inhibition of pyramidal neuron activity, and low levels of PV+ interneuron activity, with the latter fostered in part by reciprocal inhibition between CCK+ interneurons and PV+ interneurons (Caccavano & McBain, 2021; Dudok et al., 2021).

Selective loss of CCK+ interneurons in TLE could contribute to seizure generation if CCK+ interneurons are normally recruited by LEC and CA2 PN afferents to broadly suppress PN and PV+ interneuron activity during hippocampal network state transitions. CCK+ interneurons express a set of neuromodulatory receptors that may tune CCK+ interneuron activity, and thus hippocampal network excitability, based on behavioral state (Freund & Katona, 2007; Wester & McBain, 2014). The PV+ interneuron network appears relatively preserved in the epileptic hippocampus of our PILO-SE mice, and past reports of PV+ interneuron loss may in fact reflect decreases in PV expression (Wittner & Maglóczky, 2017). Thus, surviving PV+ interneurons may be disinhibited by the loss of CCK+ interneuron input and may become excessively activated by CA2 circuits, and in turn drive aberrant network synchrony in inappropriate contexts, such as network state transitions or non-REM sleep. This state-dependent property of seizure generation is suggested by the circadian oscillations in interictal discharge and seizure frequency observed in both human TLE and animal models (Baud et al., 2019, 2018; Frauscher & Gotman, 2019; Pitsch et al., 2017).

An over-activated PV+ interneuron network may promote epileptiform synchronization among PNs through mechanisms such as post-inhibitory rebound spiking (Medeiros, Cota, Oliveira, Moreira, & Moraes, 2020). As seizure activity progresses the depolarizing effects of GABAergic signaling can further drive and synchronize population firing (Ellender, Raimondo, Irkle, Lamsa, & Akerman, 2014; Hamidi & Avoli, 2015; Huberfeld et al., 2007). This model is consistent with the reports of increased interneuron activity preceding seizures (Avoli & Curtis, 2011; Muldoon et al., 2015; Neumann & Soltesz, 2017; Rich et al., 2020), including PV+ interneurons specifically (Miri, Vinck, Pant, & Cardin, 2018), and the observation that modulation of PV+ interneurons can trigger or sustain seizures (Lévesque et al., 2019; Sessolo et al., 2015; Shiri, Manseau, Lévesque, Williams, & Avoli, 2016). Additionally, PV+ interneurons may receive stronger excitatory input from disinhibited PNs, which may drive PV+ interneurons to depolarization block and thus further facilitate epileptiform discharge by PNs (Gulyás & Freund, 2015; Karlócai et al., 2014).

Like CA2 PNs, CA1 PNs in the deep *stratum pyramidale* sublayer receive strong input from PV+ interneurons (Botcher et al., 2014; Lee et al., 2014; Ribak, Seress, & Leranth, 1993; Valero et al., 2015). Thus, CA2 and deep CA1 PNs might have the greatest probability of synchronization during periods of intense neural activity. The particularly high density of PV+ interneurons in CA2, including wide-arbor PV+ basket cells and bistratified cells, could promote synchronization of CA2 PNs with nearby CA3a and CA1c neurons (Mercer, Eastlake, et al., 2012; Mercer et al., 2007). Synchronous population discharges often originate from distal CA3 and CA2 (McCloskey & Scharfman, 2011; Oliva et al., 2016; Wittner & Miles, 2007; Wong & Traub, 1983), indicating that the CA2 network has an inherent tendency towards population synchronization even in non-epileptic tissue. Thus, CCK+ interneuron impairment and other sequelae of PILO-SE may act in concert with the dense recurrent excitatory circuitry, strong entorhinal cortex input, and PV+ interneuron network in CA2 to create a powerful focus of epileptiform synchronization.

### CA2 PV+ INs appear relatively resilient in epileptic mice despite PNN disruption following PILO-SE

A previous quantitative analysis of PV+ interneurons in human TLE tissue indicated their widespread loss within the sclerotic hippocampus, including from the CA2 region (Andrioli & Arellano, 2007). Intriguingly, CA2 also showed a significant decrease in PV+ interneuron density even in non-sclerotic tissue (Andrioli & Arellano, 2007). However, when we stained for inhibitory markers to assess the survival of PV+ interneurons in PILO-SE mice we found only a modest reduction in PV+ interneuron density in CA2 (Figure 4). Moreover, the small decrease in PV+ interneuron density may reflect a loss of PV immunoreactivity, rather than a true loss of PV+ interneurons, as suggested by various studies in human TLE tissue (Wittner & Maglóczky, 2017). Additionally, our measurements of WIN-sensitive perisomatic inhibition (Figure 9) suggested that the loss of inhibition in PILO-SE mice may largely be explained by a loss or functional impairment of CCK+ INs.

The CA2 subfield contains a unique pyramidal neuron-associated PNN that has been implicated in the regulation of inhibitory synaptic plasticity at the PV+ interneuron synapses onto CA2 pyramidal neurons (Carstens et al., 2021, 2016; Domínguez et al., 2019). Additionally, PNNs are negatively regulated by CA2 PN activity levels, such that prolonged chemogenetic activation of CA2 PNs led to a decrease in PNN deposition within the CA2 region (Carstens et al., 2021). Here, we report decreased staining for CA2 PNNs in tissue from epileptic mice (Figure 5), which may reflect increased levels of CA2 PN activity leading to a downregulation of PNN deposition following PILO-SE. The CA2 PNN may be additionally impacted by seizure-associated degradation of PNN components (Carstens et al., 2021; Dubey et al., 2017; McRae et al., 2012; Rankin-gee et al., 2015). Intriguingly, recent data indicates that the CA2 PNN is decreased in the *Kcna1_-/-_* genetic model of epilepsy in mice (Carstens et al., 2021), which also presents with a pattern of mesial temporal sclerosis-like damage (Wenzel et al., 2007) suggesting similar dysregulation of extracellular matrix components across epilepsy models.

It is important to note that although our immunohistochemical measures reveal a clear decrease in the CA2 PNN, our electrophysiological data do not suggest a substantial disruption of perisomatic inhibition mediated by PV+ INs. It is possible that, despite recent evidence indicating that the PNN ensheathes PV+ perisomatic synapses onto CA2 PNs (Domínguez et al., 2019), the PNN does not contribute to the baseline magnitude of perisomatic inhibition meditated by PV+ basket cells. Rather, the PNN may be specifically involved in orchestrating inhibitory synaptic plasticity in CA2, notably the delta opioid receptor-mediated inhibitory long-term depression linked to social memory in mice (Carstens et al., 2021, 2016; Cope et al., 2021; Domínguez et al., 2019; Leroy et al., 2021; Rey et al., 2022).

## METHODS

### I. Animals

All procedures were performed in accordance with the Columbia University Institutional Animal Care and Use Committees (IACUC). Adult male and female mice (8-12 weeks-old) were housed in a temperature and humidity-controlled environment with a 12-hour light/dark cycle with food and water provided *ad libitum*. All mice were F1 generation hybrids resulting from a cross between *Amigo2*-Cre+/- mice (maintained in a C57BL/6J background, RRID:IMSR_JAX:030215) and 129S1/SvlmJ mice (Jackson Laboratory stock #002448, RRID:IMSR_JAX:002448). Genotyping was performed using tail tip samples sent to GeneTyper (Columbia University), and both Cre +/- and Cre -/- animals were used in this study.

### II. Pilocarpine-induced status epilepticus

All drugs were administered intraperitoneally (i.p.). Mice were first administered methylatropine bromide (5 mg/kg, i.p.) to suppress peripheral cholinergic activation from pilocarpine hydrochloride. Pilocarpine was administered 30 mins later (350 mg/kg, i.p.) and mice were closely and continually monitored for behavioral indicators of seizures. The onset of SE typically occurred between 30 and 60 mins following pilocarpine treatment and was defined as a convulsive seizure (stage 3, 4, or 5 on the Racine seizure scale (Racine, 1972)) that lasted continually for at least 5 mins without resuming normal behavior (e.g. grooming) for several hours. Diazepam (5 mg/kg, i.p.) was administered 1 hour after SE onset to curtail seizures. In all cases diazepam was followed 20 mins later by levetiracetam (100 mg/kg, i.p.). Control mice were given an identical timecourse of drug treatment, except they were not administered pilocarpine. Thus, control animals received methylatropine bromide, then after 150 mins were administered diazepam and at 170 mins were administered levetiracetam. Immediately after levetiracetam mice were transferred to a heated and humidified ThermoCare veterinary intensive care unit, where they were closely monitored, provided with dietary supplements in the form of DietGel Recovery (Clear H2O #72-06-5022), and given subcutaneous hydration twice daily until they showed normal locomotion and feeding (typically within 1-3 days). Control mice were returned to standard housing cages immediately following the drug administration protocol and were not given dietary supplements because they usually do not ingest them (Scharfman, unpublished observations). After recovery all mice were kept in standard group housing, except where aggression between cage-mates was observed in which cases aggressors were removed. Taking into account all mice used in this study and a previous series of experiments (Whitebirch et al., 2022), pilocarpine treatment resulted in three outcomes: acute mortality during SE due to a generalized tonic-clonic seizure leading to tonic hindlimb extension and death (114 of 411 mice, 27.7%), minor convulsive seizure activity (stage 3 seizures or below the Racine seizure scale (Racine, 1972)) without convulsive SE (133 of 411 mice, 32.4%), or convulsive SE (164 of 411 mice, 39.9%). Video-EEG recordings were not performed during induction of PILO-SE, and determination of convulsive SE was performed using established behavioral indicators including head nodding, facial automatisms and gnawing, continual bodily tremor, forelimb clonus, and periodic convulsive seizures accompanied by rearing and falling (Cavalheiro, Santos, & Priel, 1996; Lévesque, Avoli, & Bernard, 2016; Turski et al., 1984).

### III. Video-Electroencephalogram Recordings

Video-Electroencephalogram (video-EEG) recordings were performed in a subset of mice to capture spontaneous recurring seizures, the defining feature of experimentally-induced epilepsy. Surgeries to implant subdural screw electrodes (0.10” length stainless steel screws, Pinnacle Technology #8209) were performed at least four weeks after pilocarpine treatment. For surgery, carprofen was given as an analgesic agent (5 mg/kg, subcutaneous, s.c.), and postoperative analgesia was carprofen-supplemented food (either 60 mL of MediGel CPF-74-05-5022 food gel, or Rodent MDs #MD150-2 2 mg carprofen tablets, each estimated to be a dose of approximately 5 mg/kg over 24 hours of consumption). Before the commencement of surgery animals were weighed to record baseline body weight and to calculate carprofen dose. Anesthesia was achieved with isoflurane (Covetrus, NDC: 11695-6777-2), with an induction concentration of 4% and a maintenance concentration of approximately 2%. Surgical procedures were performed under sterile conditions as approved by the Columbia University IACUC. The stereotactic coordinates used were measured in millimeters relative to Bregma (for anterior-posterior and medial-lateral coordinates, A-P and M-L). Subdural screw electrodes were implanted into craniotomies located over the left and right dorsal hippocampi (A-P -1.80, M-L ± 1.30), the left frontal cortex (A-P -0.30, M-L -1.50), and the right occipital cortex (A-P -3.50, M-L +2.00). An additional two screws were placed over the right olfactory bulb (A-P +2.30, M-L +1.80) and the left cerebellum (A-P -1.5 from Lambda, M-L -0.5 from Bregma) for ground and reference electrodes, respectively. All screws were wired to an 8-pin connector (prepared from a larger block, Digi-Key ED90266-ND) and secured using dental cement (C&B Metabond #S380). Following surgery mice were individually housed and allowed to recover for a minimum of one week before video-EEG recordings were initiated. Video-EEG recording was performed with equipment from Pinnacle Technology (#8400-K1 four-channel tethered EEG system). A preamplifier (Pinnacle Technology #8400-SE4) was connected to the 8-pin connector and wired to the data conditioning and acquisition system (Pinnacle Technology #8401-HS) via a multichannel commutator (Pinnacle Technology #8408) that allowed for uninhibited movement. During long-term continuous recording sessions mice were housed in circular cages with *ad libitum* food and water and synchronous video was recorded using a box camera and infrared light source (Pinnacle Technology #9000-K10). Video-EEG recordings were acquired and reviewed using Pinnacle Technology Sirenia software (version 2.2.2). The time-frequency representation of seizure power (presented in Figure 1F) was generated in MATLAB using the cwt command, which performs a continuous 1-D wavelet transform and outputs a magnitude color scale that represents the L1 normalized power of the frequency components of the input signal.

### IV. Slice electrophysiology

#### 1. Hippocampal slice preparation

*Ex vivo* electrophysiology was performed in the chronic phase of PILO-induced epilepsy, approximately 6 weeks after status epilepticus, when mice were 14 – 18 weeks old. Acute hippocampal slices were prepared using artificial cerebrospinal fluid (ACSF) and sucrose-substituted ACSF (referred to as sucrose solution). ACSF and sucrose solutions were made with purified water that had been filtered through a 0.22 µm filter. ACSF contained (concentration expressed as millimolar, mM): 22.5 glucose, 125 NaCl, 25 NaHCO_3_, 2.5 KCl, 1.25 NaH_2_PO_4_, 3 sodium pyruvate, 1 ascorbic acid, 2 CaCl_2_, and 1 MgCl_2_. Sucrose solution used for slice preparation contained, in mM: 195 sucrose, 10 glucose, 25 NaHCO_3_, 2.5 KCl, 1.25 NaH_2_PO_4_, 2 sodium pyruvate, 0.5 CaCl_2_, 7 MgCl_2_. ACSF and sucrose cutting solution were prepared fresh before each experiment and the osmolarity was consistently measured to be between 315 and 325 mOsm. Sucrose solution was chilled on ice and bubbled with carbogen gas (95% O_2_ / 5% CO_2_) for at least 30 minutes before slice preparation. A recovery beaker was prepared with a 50:50 mixture of ACSF and sucrose solution and warmed to approximately 32° C. Mice were deeply anesthetized by isoflurane inhalation and immediate incisions were made to sever the diaphragm and access the heart. Transcardial perfusion was performed with ice-cold carbogenated sucrose cutting solution for approximately 30 – 45 seconds. The mouse was decapitated and the brain quickly removed, at which point the hippocampi were dissected free, placed into an agar block, and secured to a vibratome slicing platform with cyanoacrylate adhesive. Hippocampal slices were cut at a thickness of 400 µm, parallel to the transverse plane. Slices were collected from the dorsal and intermediate hippocampus, as CA2 circuits have primarily been characterized within the dorsal hippocampus (Chevaleyre & Siegelbaum, 2010; Hitti & Siegelbaum, 2014; Kohara et al., 2014). Slices were transferred to the warmed recovery beaker and allowed to recover for 30 minutes, after which the beaker was allowed to come to room temperature and left for an additional 90 minutes.

#### 2. Whole-cell recordings

Recording and stimulation pipettes were prepared from borosilicate glass capillaries using a heated-filament puller programmed to produce pipettes with a resistance of approximately 3 – 5 MOhm. Stimulation pipettes were filled with 1 M NaCl. Recording pipettes were filled with intracellular solution composed of, in mM: 135 potassium gluconate (C_6_H_11_KO_7_), 5 KCl, 0.2 EGTA (C_14_H_24_N_2_O_10_), 10 HEPES (C_8_H_18_N_2_O_4_S), 2 NaCl, 5 MgATP (C_10_H_16_N_5_O_13_P_3_ · Mg^2+^), 0.4 Na_2_GTP (C_10_H_16_N_5_O_14_P_3_ · Na^+^), 10 Na_2_ phosphocreatine (C_4_H_8_N_3_O_5_PNa_2_ · H_2_O), and biocytin (0.2% by weight). Intracellular solution was prepared on ice, with the osmolarity was adjusted to approximately 295 mOsm and the pH titrated to approximately 7.2. Hippocampal slices were individually transferred to the recording chamber of an electrophysiology station where ACSF, warmed to 32° C and continually bubbled with carbogen, was perfused through the chamber at a rate of approximately 1 – 3 mL/min. Whole-cell recordings from CA2 PNs was accomplished primarily through a “blind patching” approach, using the end of the *stratum lucidum* (SL) as an anatomical landmark. Cells were left unperturbed for several minutes before any recordings to ensure the membrane potential and seal were stable, and cells with a resting membrane potential more depolarized than -50 mV or a series resistance greater than 25 MOhm were discarded. The resting membrane potential was measured by recording at least one minute in the current clamp configuration, without any injected current, and averaging the membrane potential. Current was then applied as needed to hold cells at -70 mV. As previously reported (Chevaleyre & Siegelbaum, 2010; Hitti & Siegelbaum, 2014; Kohara et al., 2014; Whitebirch et al., 2022), CA2 PNs were identifiable based upon key intrinsic physiological properties including a relatively small input resistance, delayed action potential firing preceded by a gradual slow depolarizing ramp upon depolarizing current injection, a relatively small-amplitude voltage sag upon hyperpolarization, a relatively high rheobase current, and a large membrane capacitance. Analysis of intrinsic properties was performed in MATLAB and AxoGraph. Cell identity was confirmed post-hoc based on cellular morphology as revealed by biocytin and colocalization with known CA2 markers PCP4, STEP, or RGS14 (Hitti & Siegelbaum, 2014).

#### 4. Synaptic responses and pharmacology

Electrical stimulation was delivered as a single 0.2 millisecond (ms) pulse (Digitimer Ltd. Constant voltage isolated stimulator, model DS2A-Mk.II), ranging in intensity from 8 V to 80 V (in 8 V increments). For Figure 2, paired pulse stimulation consisted of two electrical stimuli separated by 50 ms. Postsynaptic potentials were measured from a holding potential of -70 mV. Fast and slow hyperpolarizing phases in Figure 3 were assessed visually for each cell: the inflection point between the fast and the slow phases of hyperpolarization typically occurred at approximately 70 ms after stimulation, and the negative peak was measured both before and after the inflection point. In experiments where inhibitory postsynaptic potentials were pharmacologically isolated, excitatory transmission was blocked with 50 µM D-AP5 (Cayman Chemical #14539) and 25 µM CNQX (Cayman Chemical #14618). The CB1 receptor agonist WIN55,212-2 (WIN, Tocris #1038) was applied at a concentration of 1 µM as indicated. Throughout the course of the experiment, stimulation was applied once every 15 seconds, and every four evoked IPSPs were averaged to produce a measurement of the IPSP at every minute. In WIN experiments the baseline IPSP was measured as the mean amplitude from at least 6 mins of stable responses. After this baseline, WIN was added to the bath for 15 mins, and then washed out for an additional 15 mins. The post-WIN IPSP amplitudes were measured as the mean amplitude of the IPSPs from minutes 11 to 25 (with the application of WIN designated as minute 0). CNQX and WIN55,212-2 were prepared as stock solutions in DMSO, and D-AP5 was prepared as a stock solution in water. Stock solutions were stored at -20 C and aliquots were thawed and kept on ice on the day of the experiment.

### V. Immunohistochemistry

#### 1. Preparation of brain sections

For immunohistochemistry mice were placed under deep isoflurane-induced anesthesia and transcardially perfused first with 0.9% NaCl and then with 4% PFA. Brains were extracted and immersed whole in 4% PFA overnight at 4° C. The next day, brains were washed in 0.3% glycine in phosphate buffered saline (PBS) for one hour at room temperature on a shaker and were then rinsed three times briefly with PBS. Coronal sections were prepared at a thickness of 60 µm using a vibratome. Post-hoc immunohistochemistry was performed by immersing 400 µm acute hippocampal slices in 4% paraformaldehyde (PFA) at the conclusion of recordings and fixing them overnight on a shaker at 4° C. Both 60 µm brain sections and 400 µm hippocampal slices were incubated for four hours at room temperature in PBS with 0.5% Triton-X. This solution was then exchanged for PBS containing primary antibodies and 0.1% Triton-X. The following antibodies were used: rabbit anti-NPY (Abcam ab30914, RRID:AB_1566510), mouse anti-parvalbumin (Sigma-Aldrich P3088, RRID:AB_477329), rabbit anti-proCCK (Frontier Institute CCK-pro-Rb- Af350, RRID:AB_2571674), rabbit anti-PCP4 (Sigma-Aldrich HPA005792, RRID:AB_1855086), mouse anti-RGS14 IgG2a (Neuromab 75-170, RRID:AB_2179931), mouse anti-STEP IgG1 (Cell Signaling Technology 4396, RRID:AB_1904101), and mouse anti-GAD67 clone 1G10.2 (Millipore MAB5406, RRID:AB_2278725). Primary antibodies were paired with goat anti-mouse and goat anti-rabbit secondary antibodies conjugated to Alexa Fluor 488 (Thermo Fisher A-21121, RRID:AB_2535764 and Thermo Fisher A-21131, RRID:AB_2535771), Alexa Fluor 568 (Thermo Fisher A-11011, RRID:AB_143157), or Alexa Fluor 647 (Thermo Fisher A-21241, RRID:AB_2535810). Neuronal somata were visualized using the Invitrogen NeuroTrace 435/455 fluorescent Nissl stain (Thermo Fisher N21479). Perineuronal nets were stained with biotinylated *Wisteria floribunda* agglutinin (WFA, Sigma-Aldrich L1516, RRID:AB_2620171). Biocytin and biotinylated WFA were both visualized using streptavidin conjugated to Alexa Fluor 647 (Invitrogen S21374, RRID:AB_2336066). These antibodies have previously been shown to have high specificity (Hitti & Siegelbaum, 2014; Kohara et al., 2014; Meira et al., 2018).

The anatomical definition of the CA2 subfield in this study is informed by prior work from our lab and others (Chevaleyre & Siegelbaum, 2010; Hitti & Siegelbaum, 2014; Kohara et al., 2014; Meira et al., 2018; Whitebirch et al., 2022). In mice the dorsal CA2 subfield is approximately 300 – 400 µm in length, and approximately three-quarters of this length overlaps with the *stratum lucidum*. This delineation of CA2 is clearly seen in the staining patterns of established markers for CA2 pyramidal cells PCP4 (Figures 1, 3, 4), STEP (Figures 1, 2), and RGS14 (Figures 5, 6). All images were acquired with a confocal microscope (Zeiss LSM 700) and processed using FIJI software (version: 2.0.0-rc-54/1.51h).

#### 2. Fluorescence and anatomical measurements

For the normalized *stratum pyramidale* Nissl fluorescence intensity profile presented in Figure 2, the proximodistal profile was normalized by length centered on the CA2 subfield. To perform the length normalization alongside Nissl fluorescence, a second proximodistal profile was measured of CA2 marker fluorescence intensity. Many hippocampal slices across experiments were combined for this analysis, and as such the CA2 marker varied between mice and included PCP4, RSG14, WFA, or GFP (the latter from expression in *Amigo2*-Cre mice). In the cases where slices were stained for multiple CA2 markers, the marker with the clearest signal-to-noise was used to generate a proximodistal profile and define the center of the CA2 subfield. A custom MATLAB script was written to transform the full length proximodistal profiles into vectors 100 values in length, consisting of 50 values from the center of CA2 to the end of CA3, and 50 values from the center of CA2 to the end of CA1. These vectors were then normalized for the intensity of each marker by dividing every value by the maximum, such that for both CA2 markers and Nissl the maximum fluorescence intensity value was 1.0 for every slice measured. In order to normalize these values against the control samples, the control Nissl intensity vectors were averaged across mice. Then, the Nissl vectors from each PILO-SE mouse were divided by the mean control value at each of the 100 positions of the length-normalized vectors, producing the data in Figure 2E. Finally, mean normalized Nissl fluorescence values were generated for each subfield (CA3c, CA3b, CA3a, CA2b, CA2a, CA1c, CA1b, and CA1a) by first defining the bounds of the CA2 subfield based on CA2 marker fluorescence. The CA2a/CA1c border exhibits a distinct transition with little intermingling of CA2 and CA1 PNs and a sharp decline in the fluorescence intensity of CA2 PN markers (Figure 2E heatmap). The CA2b/CA3a border exhibits a more gradual transition with significant intermingling of CA3 and CA2 PNs, which is reflected by a gradual decrease in CA2 marker fluorescence (Figure 2E heatmap). Therefore, the CA2b/CA3a border was drawn at the approximate halfway point of the intermingled region, so as to balance the exclusion of CA2b neurons against the inclusion of CA3a neurons. Considering recent evidence for proximodistal heterogeneity within the CA2 region (Fernandez-lamo et al., 2019; Okamoto & Ikegaya, 2018), and our own observations of greater resilience to neurodegeneration in distal CA2, the CA2 subfield was divided into two equal halves designated as CA2a (distal, near CA1) and CA2b (proximal, near CA3). Such a subdivision of the CA2 subfield has been proposed as a means to take into account the anatomical distinctions between the proximal and distal ends of CA2, considering that neurons in the latter region do not colocalize with the DG mossy fibers (Dudek, Alexander, & Farris, 2016; Fernandez-lamo et al., 2019; Okamoto & Ikegaya, 2018). The CA3 and CA1 regions were each divided in three equal regions by length. The normalized Nissl fluorescence intensity values were averaged in each region to produce the values represented in Figure 2F.

PNN integrity was assessed by immunostaining with biotinylated WFA. The bounds of the CA2 subfield were defined based upon co-staining with PCP4. In FIJI regions of interest were drawn around the CA2 *stratum oriens* (SO), *stratum pyramidale* (SP), and *stratum radiatum* (SR) and the mean WFA fluorescence intensity measured in each region of interest. Regions of interest were drawn to exclude WFA associated with PV+ interneurons. In the dentate gyrus, regions of interest were drawn in a portion of the granule cell layer (GCL), molecular layer (ML), and hilus (HIL) (measurements were made in the upper blade of the DG). For every animal, background fluorescence intensity was subtracted from the measured WFA fluorescence intensity and measurements from multiple hippocampal sections were averaged to produce a single set of measurements from each animal. To measure WFA associated with PV+ interneurons, sections were co-stained for parvalbumin, regions of interest were drawn around several PV+ interneuron soma within the CA2 subfield visualized with PCP4, and the WFA fluorescence intensity from all measured interneurons was averaged for each animal. Parvalbumin (PV) and pro-cholecystokinin (pCCK)-expressing interneuron density was measured in hippocampal sections stained with the PV and pCCK antibodies described above, along with a stain against PCP4 or RGS14. The CA2 region was captured in a Z-stack series of images and the bounds of the CA2 subfield were drawn across all layers of the hippocampus using PCP4 or RGS14 immunofluorescence. For pCCK+ interneuron measurements, regions of interest were additionally drawn across all layers of CA3 and CA1, with each subfield divided lengthwise into approximate thirds (CA3c, CA3b, CA3a, CA1c, CA1b, CA1a). As with WFA fluorescence intensity, interneuron density measurements from multiple sections were averaged to produce a single set of measurements for each mouse. All imaging was performed with identical acquisition parameters for samples from control and PILO- SE mice.

### VI. Statistical comparisons

All statistical tests were performed in GraphPad Prism. For group comparisons, two-way ANOVA with the Geisser-Greenhouse correction and Holm-Sidak’s multiple comparisons test was used. For group comparisons with missing data points, GraphPad Prism cannot implement a two-way ANOVA and instead utilizes a mixed-effects model that uses a compound symmetry covariance matrix and which is fit using the restricted maximum likelihood (REML) approach. The Mann- Whitney U test was used for comparisons between two data sets, and the Kruskal-Wallis test with Dunn’s multiple comparisons test was used for comparisons between more than two groups. Comparison of paired datasets was performed using the Wilcoxon matched-pairs signed rank test. For immunohistochemical and anatomical data, statistical tests were performed using the number of mice as the sample size n, with measurements from multiple tissue sections averaged to yield a single set of values for each animal. In the case of electrophysiological measurements, n was defined as the number of individual neurons. In all cases the cutoff for significance was set to a P value of 0.05.

## Acknowledgements

This work was supported by National Institutes of Health grants R01NS106983 from NIH (PIs, H.E.S. and S.A.S.) and grant F31NS113466 (A.C.W.)

